# Efficient Hit-to-Lead Searching of Kinase Inhibitor Chemical Space via Computational Fragment Merging

**DOI:** 10.1101/2021.06.01.446684

**Authors:** Grigorii V. Andrianov, Wern Juin Gabriel Ong, Ilya Serebriiskii, John Karanicolas

## Abstract

In early stage drug discovery, the hit-to-lead optimization (or “hit expansion”) stage entails starting from a newly-identified active compound, and improving its potency or other properties. Traditionally this process relies on synthesizing and evaluating a series of analogs to build up structure-activity relationships. Here, we describe a computational strategy focused on kinase inhibitors, intended to expedite the process of identifying analogs with improved potency. Our protocol begins from an inhibitor of the target kinase, and generalizes the synthetic route used to access it. By searching for commercially-available replacements for the individual building blocks used to make the parent inhibitor, we compile an enumerated library of compounds that can be accessed using the same chemical transformations; these huge libraries can exceed many millions – or billions – of compounds. Because the resulting libraries are much too large for explicit virtual screening, we instead consider alternate approaches to identify the top-scoring compounds. We find that contributions from individual substituents are well-described by a pairwise additivity approximation, provided that the corresponding fragments position their shared core in precisely the same way relative to the binding site. This key insight allows us to determine which fragments are suitable for merging into a single new compounds, and which are not. Further, the use of the pairwise approximation allows interaction energies to be assigned to each compound in the library, without the need for any further structure-based modeling: interaction energies instead can be reliably estimated from the energies of the component fragments, and the reduced computational requirements allow for flexible energy minimizations that allow the kinase to respond to each substitution. We demonstrate this protocol using libraries built from six representative kinase inhibitors drawn from the literature, which target five different kinases: CDK9, CHK1, CDK2, EGFR^T790M^, and ACK1. In each example, the enumerated library includes additional analogs reported by the original study to have activity, and these analogs are successfully prioritized within the library. We envision that the insights from this work can facilitate the rapid assembly and screening of increasingly large libraries for focused hit-to-lead optimization. To enable adoption of these methods and to encourage further analyses, we disseminate the computational tools needed to deploy this protocol.

**Graphical Abstract:** 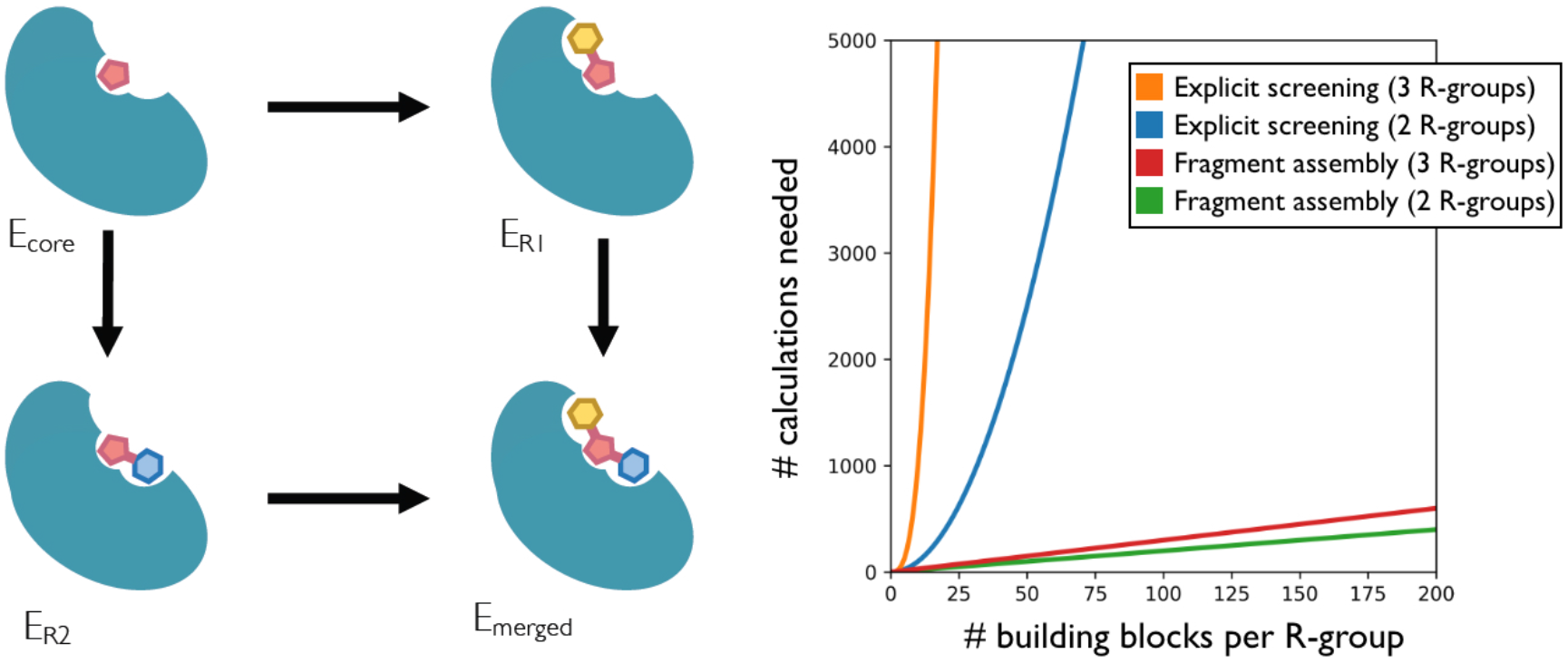

## Introduction

The practice of virtual screening has historically comprised two separate and complementary tasks, irrespective of a campaign’s particular target or goals. The first task focuses on finding active compounds (“hits”) by rapidly docking a large collection of drug-like compounds against some protein target [1]. Historically, this has most often been carried out by using fast and approximate methods for evaluating each compound comprising the ubiquitous ZINC database (a concatenation of numerous vendor’s catalogs that spanned 3-8 million diverse compounds) [2, 3]. The second task aims to take the validated screening hits (whether from computational or biochemical screens), and to optimize their activity by proposing improved analogs [4, 5]. Here, expensive free energy perturbation methods (or equivalent) are typically used, applied at the scale of about a hundred compounds in a highly focused part of chemical space [6].

In the past few years, certain chemical vendors began listing not just the contents of their shelves, but also enumerating all the compounds that can be readily synthesized from these building blocks [7]. Inclusion of these enumerated “make-on-demand” compounds led to a dramatic and immediate increase in the size of libraries for virtual screening. At the same time, by virtue of the underlying synthetic strategy, many variants of a given chemical scaffold are proffered in which a given building block can be replaced with thousands of related alternatives: thus, these new collections also afford incredibly *dense* coverage of chemical space. The availability of numerous close analogs within the screening collection allows for the possibility that the first virtual screening task may not simply deliver starting points for future hit-to-lead optimization, but rather that it might instead provide potent compounds that will not require extensive further refinement. The suggestion that potent compounds might be identified from the initial screening stage now blurs the classical sharp delineation between the “hit finding” and “hit-to-lead” tasks.

The sheer size of these enumerated collections – now at least 29 billion compounds [8] – has forced practitioners to reconsider how this chemical space should be sampled, given the impracticality of sequentially docking each of these compounds to one’s target protein even with massive computational resources and emerging GPU capabilities [9, 10]. One potential approach might be to screen a subset of the collection, with the goal of identifying “privileged” scaffolds, then subsequently re-screen each of the available compounds that elaborate this scaffold in different ways [11–13]. However, two separate studies have by now demonstrated that valuable hits would be missed this way: one cannot necessarily recognize from the scaffold alone that an elaborated analog might score well [7, 13].

For certain well-studied target classes, though, clusters of “privileged” chemotypes have already been identified. This is especially true in the case of protein kinases, where the structurally-conserved ATP-binding site has led to the repeated re-use (or inadvertent re-discovery) of a broad set of hinge-binding cores [14]. By starting from a given core and choosing different substituents, a particular core can be developed in different ways; thus, selective inhibitors for many different kinases can be built from the same core [15]. This insight further serves as the basis for synthesis of numerous kinase-focused chemical libraries [16], which harbor a limited diversity of chemotypes but are poised to be optimized for any kinase of interest.

Here, we seek to address the challenge of how one might efficiently search the narrow but very dense swath of chemical space around a given core, to optimize it for a particular target. We choose six diverse inhibitors of different kinases as starting points, and in each case generate and explore the chemical space around this inhibitor. We describe a “deconstruction-reconstruction” approach [17, 18], in which the synthetic route used to build the known inhibitor is generalized, and used to create a huge chemical library that densely samples synthetically-accessible chemical space. The resulting libraries can exceed billions of compounds, vastly exceeding the scales amenable to explicit docking even when the protein is held fully fixed [7]. We then find that by adapting strategies that underlie fragment merging [19, 20], the top-scoring compounds from the resulting library can be identified in an extremely efficient manner, while allowing for protein conformational changes necessary to recognize active analogs of the parent compounds.

## Model Systems

Protein kinases are a thoroughly established class of targets for therapeutic intervention, particularly in oncology, due in part to their central role in mediating cellular signaling [21]. Protein kinases share a highly conserved catalytic domain (**Figure 1a**), which is responsible for transferring a phosphate group from ATP to a substrate protein, and in so doing altering the recipient protein’s activity. While their inactive states can vary, the active conformation of protein kinases is essentially invariant [22–24]. In this conformation the kinase engages ATP via a specific pose, through set of specific interactions; accordingly, the ATP binding site exhibits exceedingly strong structural conservation. Most kinase inhibitors bind to this site, disrupting kinase activity by competing with ATP. The majority of these – known as “Type I” inhibitors – occupy almost exactly the same space as ATP within the binding site, and they are thus compatible with the kinase’s active conformation.

**Figure 1:**
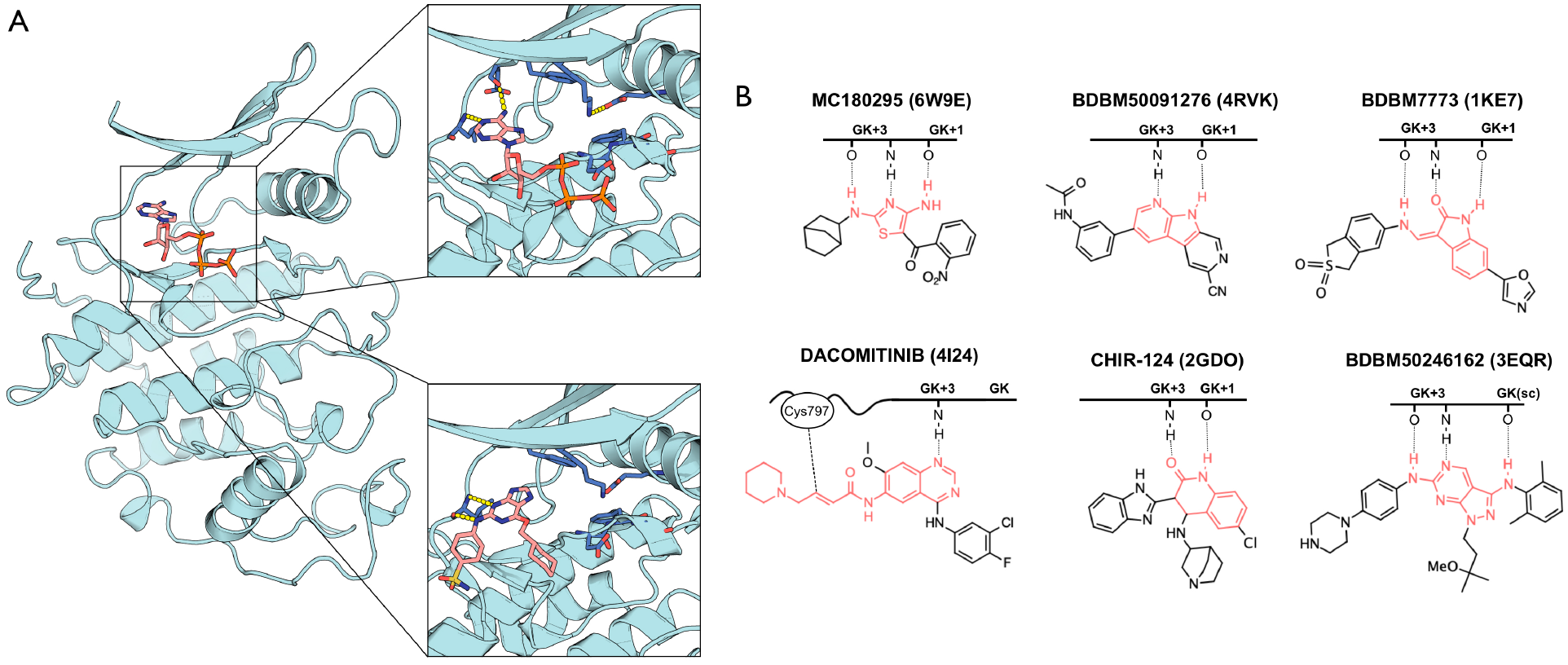
The six kinase inhibitors used as the focus for our study. (**A**) The catalytic domain of a representative protein kinase, cyclin-dependent kinase 2 (CDK2). Crystal structures in complex with ATP (*upper inset*) and in complex with a representative Type I inhibitor (*lower inset*) demonstrate the similarity of the protein conformation, and the inhibitor’s mimicry of the interactions between ATP and the kinase. (**B**) Each of the kinase inhibitors used as starting point for our study have a clear hinge-binding motif (*red*). The kinase hinge region is represented schematically, with numbering relative to the gatekeeper (GK) residue. The pattern of hydrogen bonds between each inhibitor and the kinase hinge presented here is drawn from the crystal structures of each complex (PDB IDs are provided for each complex). Each of these inhibitors engages the kinase hinge via two or three hydrogen bonds to the GK+1 and GK+3 positions, except Dacomitinib (which makes only one hydrogen bond, but forms a covalent bond to Cys797). Inhibitor BDBM50246162 also includes a hydrogen bond to the sidechain of the GK residue (threonine).

By virtue of engaging a structurally conserved binding site evolved to recognize ATP, most Type I inhibitors in fact mimic the three-dimensional used by ATP to bind the kinase. ATP’s flat adenine moiety is sandwiched between hydrophobic groups above and below, and the adenine’s edge forms specific hydrogen bond interactions with backbone groups on the “hinge” region that connects the N- and C-lobes of the kinase. The same can be said for nearly all Type I kinase inhibitors: though they rarely use adenine itself, they instead use alternate flat rings that present a comparable pattern of polar groups on their hinge-facing edge, allowing them to engage the kinase in a manner very reminiscent of ATP. Indeed, the repertoire of hinge-binding cores repeatedly used by medicinal chemists to engage kinases (mentioned earlier) form the basis for nearly all known kinase inhibitors [14], as well as for kinase-focused chemical libraries [16].

To evaluate our computational approach, we selected six diverse kinase inhibitors as starting points for study. We chose these particular examples by virtue of the crystal structure of these inhibitors having been solved in complex with the kinase (to allow retrospective analysis), a clear hinge-binding motif (**Figure 1b**), and straightforward synthetic routes. By these criteria, many more inhibitors would also qualify; we chose these six examples for study based on their chemical dissimilarity relative to one another.

Each of the six compounds selected for study inhibits its target kinase with IC_50_ better than 20 nM, through the culmination of a medicinal chemistry campaign described in each of the studies reporting these compounds. Moreover, the results of these optimization campaigns also provided at least 10 analogs of each inhibitor that have IC_50_ better than 1 μM.

For clarity we will refer to each of these inhibitors by their names in PubChem. The six inhibitors are:

1) MC180295, a selective CDK9 inhibitor built on a diaminothiazole core [25, 26] (PDB ID 6W9E)
2) BDBM50091276, a CHK1 inhibitor built on a 1,7-diazacarbazole core [27] (“compound 8” in the original paper, PDB ID 4RVK)
3) BDBM7773, a CDK2 inhibitor built on an oxindole core [28] (“compound 109” in the original paper, PDB ID 1KE7)
4) Dacomitinib (PF00299804), a covalent EGFR^T790M^ inhibitor built on a 4-aminoquinazoline core [29, 30] (PDB ID 4I24)
5) CHIR-124, a CHK1 inhibitor built on a quinolinone core [31] (“compound 11” in the original paper, PDB ID 2GDO)
6) BDBM50246162, an ACK1 (aka TNK2) inhibitor built on a pyrazolopyrimidine-3,6-diamine core [32] (“compound 2” in the original paper, PDB ID 3EQR)

Our task will be to build a chemical library by diversifying around this compound, and then to efficiently identify the best analogs from this library. Below, we describe an approach for rapidly carrying out this search using a strategy inspired by fragment merging. In the *Results* section, which follows the description of our computational approach, we will detail the specifics of applying this approach using these six representative cases as concrete examples.

## Computational Approach

### Step I: Retrosynthetic analysis

Starting from the known inhibitor, we use retrosynthetic analysis to generalize the synthesis of this compound (**Figure 2**, *Step I*). Computational methods for synthetic route planning have made rapid and dramatic advances in recent years [33–39], and provide a natural tool to carry out this step. That said, for the current study we have drawn inhibitors from the literature, and thus we have access to already-validated synthetic routes to access these compounds. By re-using the previously-described syntheses, we expect that many of the compounds described through the reported structure-activity relationships will also be present in our designed library, facilitating benchmarking in this analysis. For this reason, we have elected in the present study to simply re-use the synthetic routes reported in the literature for each starting compound.

**Figure 2:**
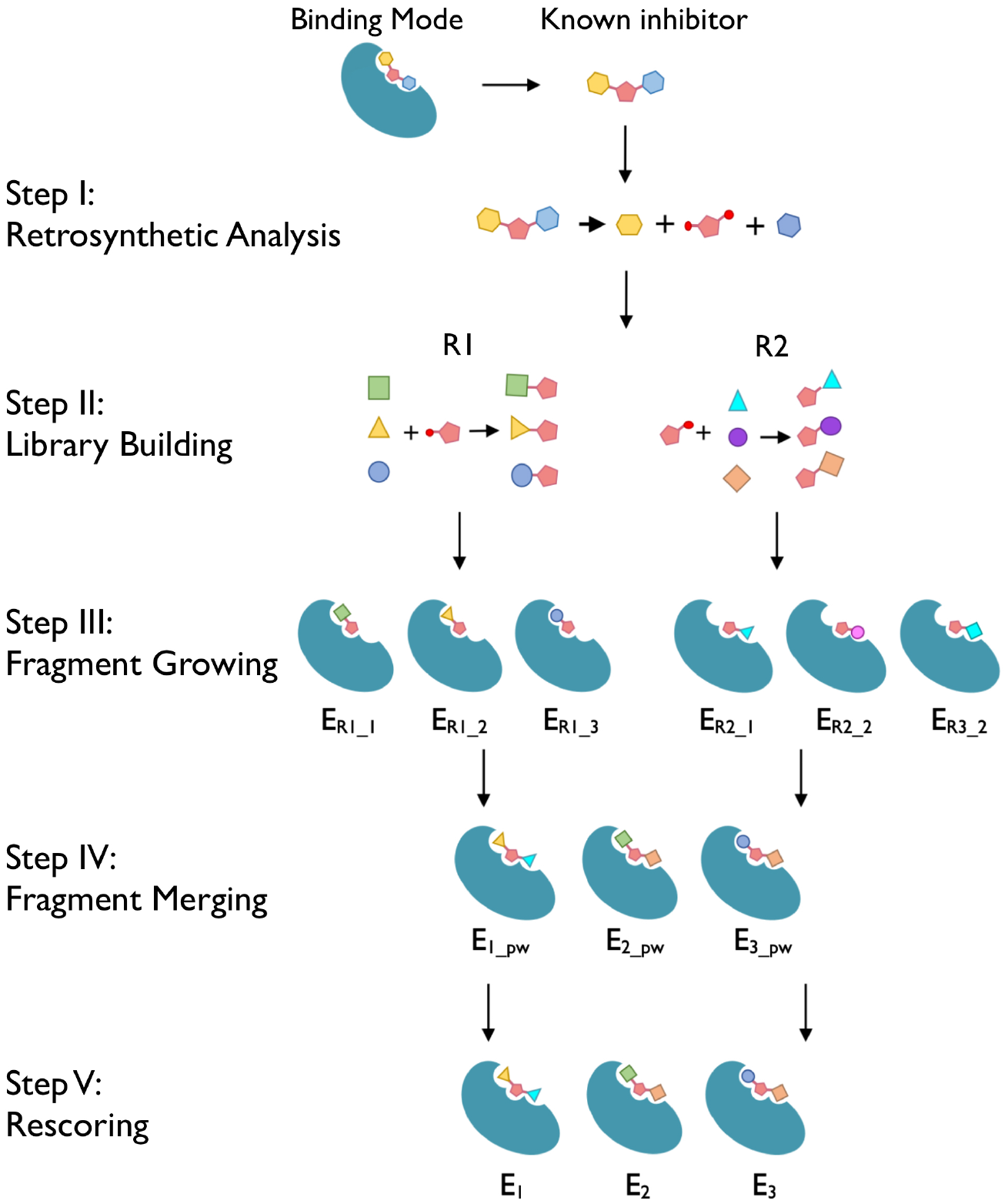
Computational fragment merging strategy for efficient hit-to-lead optimization of kinase inhibitors. Summary of our computational fragment merging approach. *Step I:* Starting from a known inhibitor, we use retrosynthetic analysis and its crystal structure in complex with the kinase to decompose the inhibitor into its hinge-binding motif (*pink pentagon*) and its side chains (*blue and yellow hexagons*). *Step II:* Diversity at the side chains is introduced by identifying alternate (commercially-available) building blocks that can be used in place of the original reactants. These are then joined to the hinge-binding scaffold to form newly-elaborated hinge-binding fragments. *Step III:* Each of the elaborated hinge-binding fragments are separately aligned into the kinase’s ATP-binding site, and refined via energy minimization. *Step IV:* Fragments elaborated at different positions can be merged if the shared core is positioned in precisely the same orientation. Given the shared positioning of the core, the interaction energy of the merged compound can be estimated from the interaction energies of the component fragments (assuming pairwise additivity). *Step V:* The merged inhibitor is then re-refined via energy minimization, and the interaction energy calculated explicitly (without assuming pairwise additivity).

### Step II: Library building

The goal of our approach is not to search alternate potential hinge-binding cores, but rather to exhaustively sample elaborations of the current core. The hinge-binding motif is almost always located in the middle of the inhibitor (due to the three-dimensional architecture of the kinase binding site), and so we began by (manually) identifying this motif and keeping it fixed.

For each of the building blocks appended onto the hinge-binding motif as part of the original reaction, we sought to retain the functional group(s) needed for the chemical transformation and diversify the rest of the building block. To do so, we first wrote the simplest unsubstituted version of each building block as a SMARTS string (**Table S1**). Briefly, a SMARTS string represents a molecular pattern [40]; in our case, we use these patterns to encode an abstraction of the starting building block. Thus, the SMARTS string can serve as a query to identify other compounds that could be used in the original reaction, as a substitute for the original building block. Using each SMARTS string as a query, we then searched PubChem to identify alternate building blocks from commercial vendors that could be used for this reaction (**Figure 2**, *Step II*), with a series of additional chemical filters on the allowed hits (**Table S2**) and restrictions on the chemical vendors to include in the search (**Table S3**).

For each of the candidate building blocks, we then used the “ChemicalReaction” functionality in RDKit [41] to assemble the product of the corresponding chemical transformation. To build the complete collection of inhibitors that can be assembled from these building blocks, we wrote SMARTS strings that encode all steps to generate the final (complete) inhibitor from an arbitrary set of building blocks. However, because we planned primarily to evaluate each substituent separately (as will be described below), we also wrote SMARTS strings to generate a corresponding fragment (i.e., one new substituent added onto the hinge-binding core) from each building block (**Table S1**).

### Step III: Fragment growing

By diversifying multiple building blocks from the original reaction and exhaustively recombining them, one can thus generate very large libraries of new compounds built on the original hinge-binding motif, that can (in principle) be accessed using the original synthetic route. Depending on the size of the library, though, it can be prohibitive to explicitly screen these.

The positioning of the hinge-binding motif within the kinase active site is typically conserved, because these motifs form hydrogen bonds to specific backbone groups on the kinase. To provide a compatible binding mode with the kinase active site, the enumerated compounds in this chemical library must therefore be accommodated in the kinase binding site without disrupting the positioning of the hinge-binding motif.

Extending the hinge-binding motif by addition of a new group is strongly reminiscent of the “growing” approach in fragment-based drug discovery (FBDD) (or fragment-based lead discovery, FBLD) [19, 20]. As with fragment growing, our task is to entails adding new groups onto the starting hit, to form additional adventitious interactions with the protein target. Indeed, the first fragment-derived drug to reach the clinic was the kinase inhibitor vemurafenib (targeting BRAF V600E) [42], designed by growing the initial hinge-binding fragment 7-azaindole [43]. Some computational approaches for fragment growing have used docking or free energy perturbation methods to guide which expansions to make and test [44–46]; in the case of the kinase active site, however, structural conservation of the hinge-binding motif greatly restricts potential poses of the elaborated compound.

Rather than allow extensive conformational searching, then, we placed each elaborated compound into the kinase active site by alignment of the hinge-binding motif (**Figure 2**, *Step III*). For each potential elaboration of the hinge-binding motif accessible using the building blocks from the previous step, we generated low-energy conformations using the OMEGA software [47]. Each of these conformers have a shared substructure with the original inhibitor (the hinge-binding motif), allowing them to be individually aligned into the kinase active site (by overlay of the hinge-binding motif, using RDKit [41]). Each protein-ligand complex was subjected to energy minimization using the Rosetta scoring function [48, 49], and the lowest-energy conformer that included the expected hinge-binding hydrogen bond pattern was carried forward.

### Step IV: Fragment merging

Having separately optimized individual substituents in conjunction with the hinge-binding motif, our next goal was to combine these into a single chemical entity. This task is encountered elsewhere in a separate class of fragment-based drug discovery, fragment merging. Broadly speaking, merging can be used when two different fragments both include a shared functional group that engages the protein the same way, but the two have substituents at different positions; in this case the substituents from each of the two fragments can be joined onto the shared core, and the substituents’ interactions with the protein can be preserved [19, 20].

By construction, our protocol brings us to the point of having variations on a shared core (the hinge-binding motif) with substituents at different positions. Given their shared cores, pairs of fragments can naturally be merged to yield a new candidate kinase inhibitor (**Figure 2**, *Step IV*). All potential pairs are exhaustively considered, and thus the effective chemical space spanned by this approach can be very large. Fragments can best be merged if the shared core occupies precisely the same pose in both fragments; if either substituent makes interactions with the protein that require changes to the positioning of the shared core, they are fundamentally incompatible with one another. For pairs of fragments that elaborate the hinge-binding motif at different positions, the RMSD for the shared core (hinge-binding motif) can be used as a measure of their compatibility. As we will demonstrate later, fragment pairs with low RMSD yield interaction energies that are readily predicted from the sums of their component parts.

This fragment merging approach also naturally provides the expected pose for this new inhibitor, by simply retaining the coordinates of the shared core, and concatenating together the substituents’ coordinates from the corresponding fragments. Importantly, since this fragment merging approach combines the structures of substituents in each one’s preferred geometry, one might assume additivity of their interactions. This implies that the interaction energy of the newly-constructed compound with the kinase can be estimated from the interaction energies of the component fragments. This will be explored further using the real examples to follow, but the assumption of energetic additivity between substituents allows the top-scoring compounds to be identified at this merging stage, without the need to explicitly refine all models.

### Step V: Selecting top-scoring inhibitors

Finally, we re-refine and re-score the top-scoring compounds from the preceding step, by energy minimization using Rosetta [48, 49] (**Figure 2**, *Step V*). This step serves to ensure that the selected substituents on the shared hinge-binding core are indeed compatible with one another, both structurally and energetically. Slight differences in the orientation of the shared core can lead to slight deviations from additivity for the substituents, and thus refining and re-ranking them explicitly ensures that the top-scoring new candidate inhibitors are identified for further characterization.

## Results

### Library building from generalized reaction schemes

For each of the six inhibitors to be considered (**Figure 1**), we used as a starting point the synthetic scheme used to the prepare each compound. As noted earlier, computational methods for synthetic route planning could be used for this step; however, re-using the reported path makes it more likely that analogs reported in each campaign will also be present in our computational library, facilitating benchmarking for this study.

As is typical for preparation of kinase inhibitors, the synthetic schemes are designed to elaborate a particular hinge-binding core with many diverse substituents. The building blocks that contribute groups outside the core are ideally used late in the synthetic route, so that common intermediates may be re-used.

Using the approach described earlier, we generalized each reaction and searched for building blocks matching the patterns required for each chemical transformation. The libraries resulting from enumeration of commercially-available building blocks are, unsurprisingly, astoundingly large (**Figure 3**). In generalizing the synthetic route used to access MC180295, for example, our search revealed about 5,000 2-bromoacetophenones, and 750,000 primary amines: enumerating all pairs leads to almost 4 *billion* diaminothiazoles that could be accessed using this synthetic route. In fact, many more potential inhibitors can be reached though this path, through the trivial synthesis of additional 2-bromoacetophenone building blocks; for this first study, though, we limit our search to the lower bound of libraries that can be immediately accessed using available starting materials.

**Figure 3:**
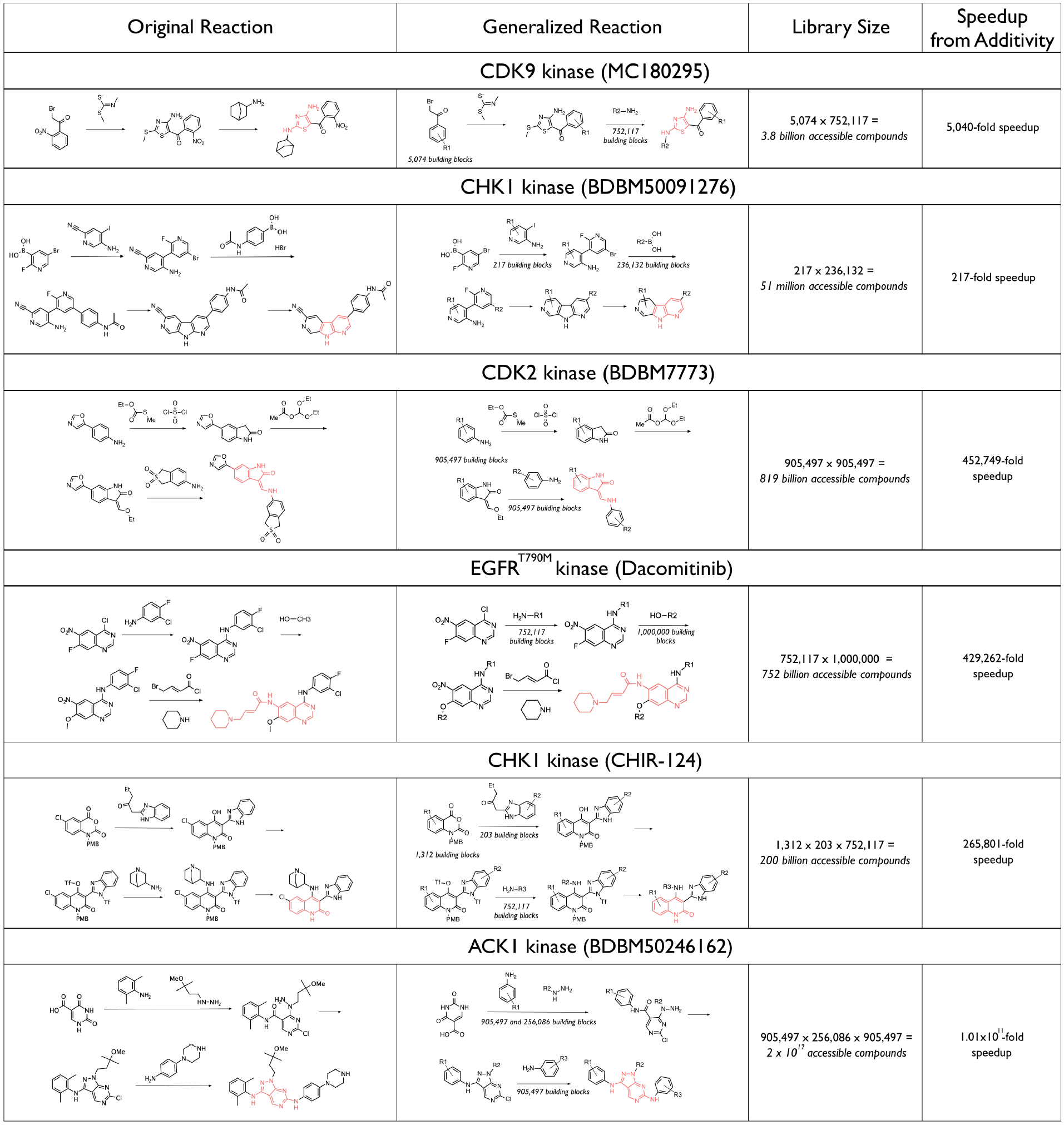
Library building from generalized reaction schemes. *First column:* The reported synthetic routes for the five representative kinase inhibitors are shown, with the hinge-binding core highlighted in *red*. *Second column:* Generalizing each reaction by allowing for commercially-available building blocks with diversified substituents enables generation of new compounds that can be accessed through the same synthetic route. *Third column:* Enumerating the chemical space accessible through this strategy demonstrates the magnitude of libraries compiled in this manner. *Fourth column:* By screening individual fragments and then assuming pairwise additivity to estimate the energy of each compound in the enumerated library, the computational demands of screening these libraries can be dramatically reduced.

As expected, the number of available building blocks at each stage is determined by complexity and popularity of the template: generic primary amines and alcohols, boronic acids, and arylamine (found in the generalized syntheses of MC180295, BDBM50091276, Dacominitib, and BDBM7773 respectively) yield many hundreds of thousands of matching building blocks. By contrast, more specific patterns or those that correspond to obscure starting materials may yield only hundreds of potential alternatives. Naturally, the number of building blocks available for diversification also strongly influences the resulting library size: in the case of BDBM50246162, appending three different substituents from readily-available building blocks results in a much larger chemical space in which to search (2 x 10^17^ compounds).

For each reaction, we first searched the newly-generated compounds to determine whether the known inhibitor was present in the enumerated library. Essentially, this test simply evaluates whether the required building blocks for this compound were identified from PubChem: in all six cases, this was confirmed to be the case. Moreover, each library also included many of the analogs described by the original authors of each study, allowing for further benchmarking.

### Assembling models of bound inhibitors from fragments

Before testing our fragment-based computational approach in a screening context, we first sought to examine whether building up the structure of the known inhibitor from fragments would yield the correct (experimentally determined) pose. For each of the six case studies, we therefore began by applying our computational approach using the fragments that yield the known inhibitor.

As described earlier, low-energy conformers are built from each fragment. Each conformer is then separately aligned into the kinase active site using the hinge-binding motif, subjected to energy minimization, and the fragment with the best interaction energy is carried forward (**Figure 2**, *Step III*). Using fragments from each of the known inhibitors, we found in all cases that the elaborated functional group adopted a conformation very similar to that observed in the crystal structure of the full inhibitor (**Figure 4a**). Further, we were gratified to see in all cases that the resulting poses retained the intended hinge-binding motif in the appropriate conformation: should either pose position this core in an alternate position, it would not be possible to merge the two fragments.

**Figure 4:**
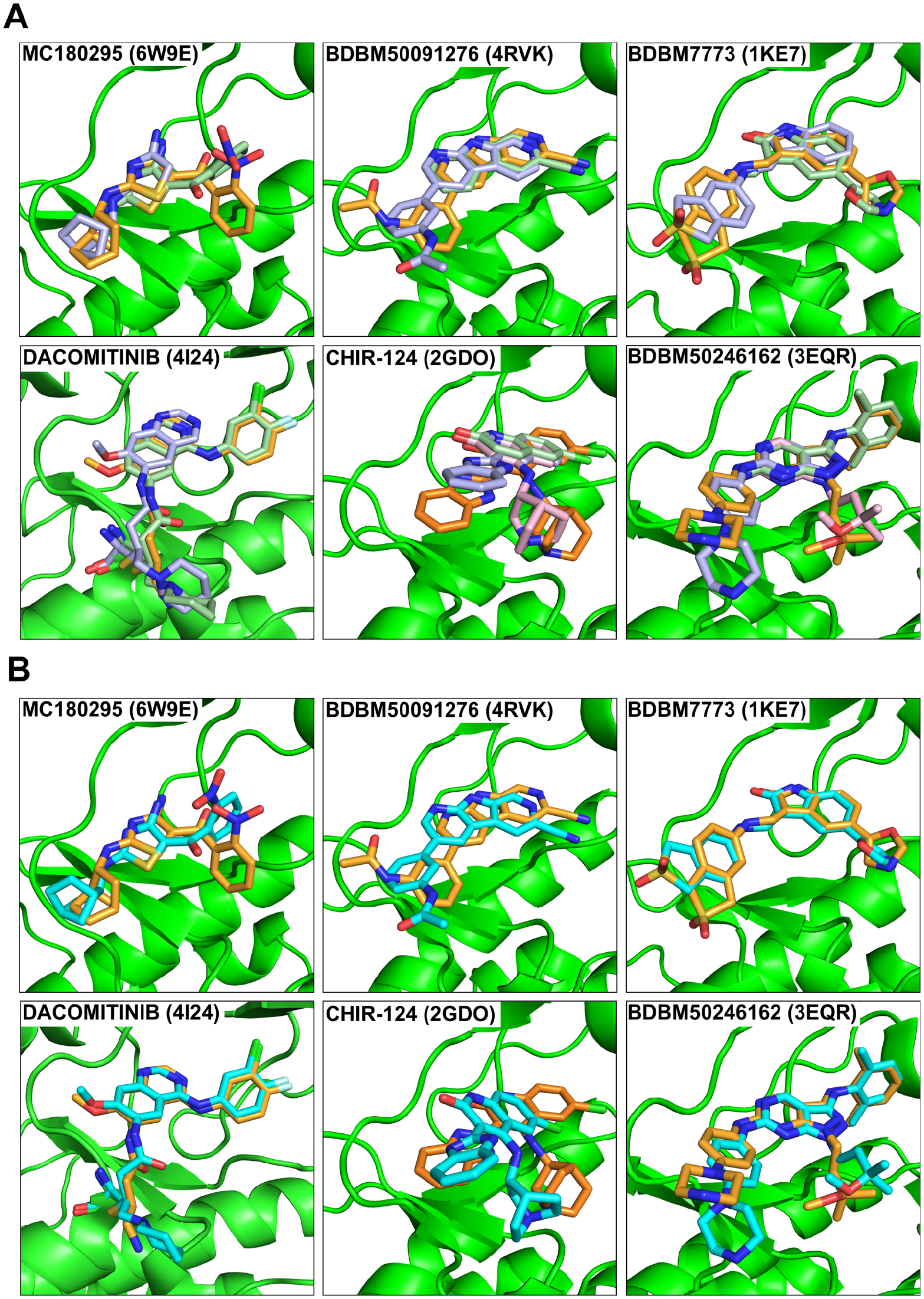
Recapitulation of binding mode for known inhibitors. (**A**) For each of the five known inhibitors, we began from fragments corresponding to the substituents in the known inhibitor (i.e., each fragment comprises the hinge-binding core and one of its side chains). Models were prepared for each fragment in complex with the cognate kinase, and then refined via energy minimization. The resulting models of the fragments (*slate and light green*) position the side chains in close agreement with their positions in the crystal structure of the complete inhibitor (*orange*). Moreover, the positioning of the hinge-binding core in both fragments is closely overlaid, allowing the fragments to be merge. (**B**) Merging the fragments and refining the resulting model by energy minimization led to models of the complete inhibitor (*cyan*) that closely match the crystal structures of these compounds (*orange*).

This was an important and reassuring result, because it implied that – at least for these potent and well-optimized inhibitors – there is no “tug-of-war” between the two sides of the inhibitor. Rather, the individual contributions of the substituents added to the central core are also locally optimal, because their interactions with the protein are the same irrespective of whether other substituents are present on this core.

We then merged each of the cognate fragment pairings (**Figure 2**, *Step IV*), and re-minimized this model of the full inhibitor (**Figure 2**, *Step V*). Unsurprisingly, given that the initial model had been built from complementary fragments with a closely shared core, minimization of the complete inhibitor did not result in drastic changes to the pose. Moreover, since the fragments were already in good agreement with the corresponding parts of the inhibitors’ crystal structures, the resulting models of the complete inhibitors also closely matched the corresponding experimentally determined structures (**Figure 4b**).

### Pairwise additivity of interaction energies

When extending this fragment-merging strategy to a screening context, our intention is to locally optimize the component fragments (**Figure 2**, *Step III*), then merge them only if the shared core aligns in both cases. A fragment that shifts the positioning of the shared core relative to the starting orientation might be used in an inhibitor, but it can only be credibly merged with another fragment that also shifts the shared core in precisely the same way.

A consequence of merging fragments with structurally-shared cores is that their interactions with the kinase are expected to be energetically additive. Non-additivity between pairs of chemical substitutions can arise for a number of reasons: primarily if the preferred conformation for the two substituents clash with one another, or each requires the protein to adapt in a way that is inconsistent with the other, or due to changes in solvent structure, or if they lead to mutually-incompatible conformation of their shared cores [50–53]. In each of our testcases, substituents are added at vectors pointing in opposing directions, making it unlikely that the substituent at one position (or its effect on the kinase conformation) would be felt at the other position. Individual water molecules are not explicitly modeled in our energy function, eliminating this potential source of non-additivity. Thus, we hypothesized that the interaction energy of the kinase inhibitors comprising our libraries could be predicted from the energies from the fragments, provided that the location of the shared core is the same in both fragments.

To test this hypothesis, we sought to evaluate the expected interaction for a full inhibitor from its fragments (given the assumption of pairwise additivity), and then explicitly assembled this inhibitor and explicitly evaluate its interaction energy (after minimization of the full inhibitor). Because this experiment requires explicit minimization of each candidate inhibitor (including the protein degrees of freedom), it cannot be carried out to completion using the huge libraries of accessible compounds described earlier (**Figure 3**). Instead, we randomly selected a subset of fragments for each building block, and exhaustively enumerated smaller libraries of accessible compounds (typically several hundred thousand compounds) (**Table S4**).

We built models corresponding to each of the fragments in this library as described earlier (**Figure 2**, *Step III*), minimized them in complex with the kinase, and determined their interaction energies. Based on a standard chemical double-mutant cycle [54] (**Figure 5a**), albeit here using interaction energies from Rosetta rather than true binding free energies, we evaluated the interaction energy expected for each compound in our library. We then merged each of the cognate fragment pairings (**Figure 2**, *Step IV*), and minimized this model of the full inhibitor (**Figure 2**, *Step V*). Unsurprisingly, the distribution of atomic contacts within sub-pockets of the binding site by the merged inhibitor (as defined by KinFragLib [55]) matched the union of the component fragments (**Figure S1**), supporting the expectation that assembling the inhibitors in place typically does not dramatically alter the bound pose.

**Figure 5:**
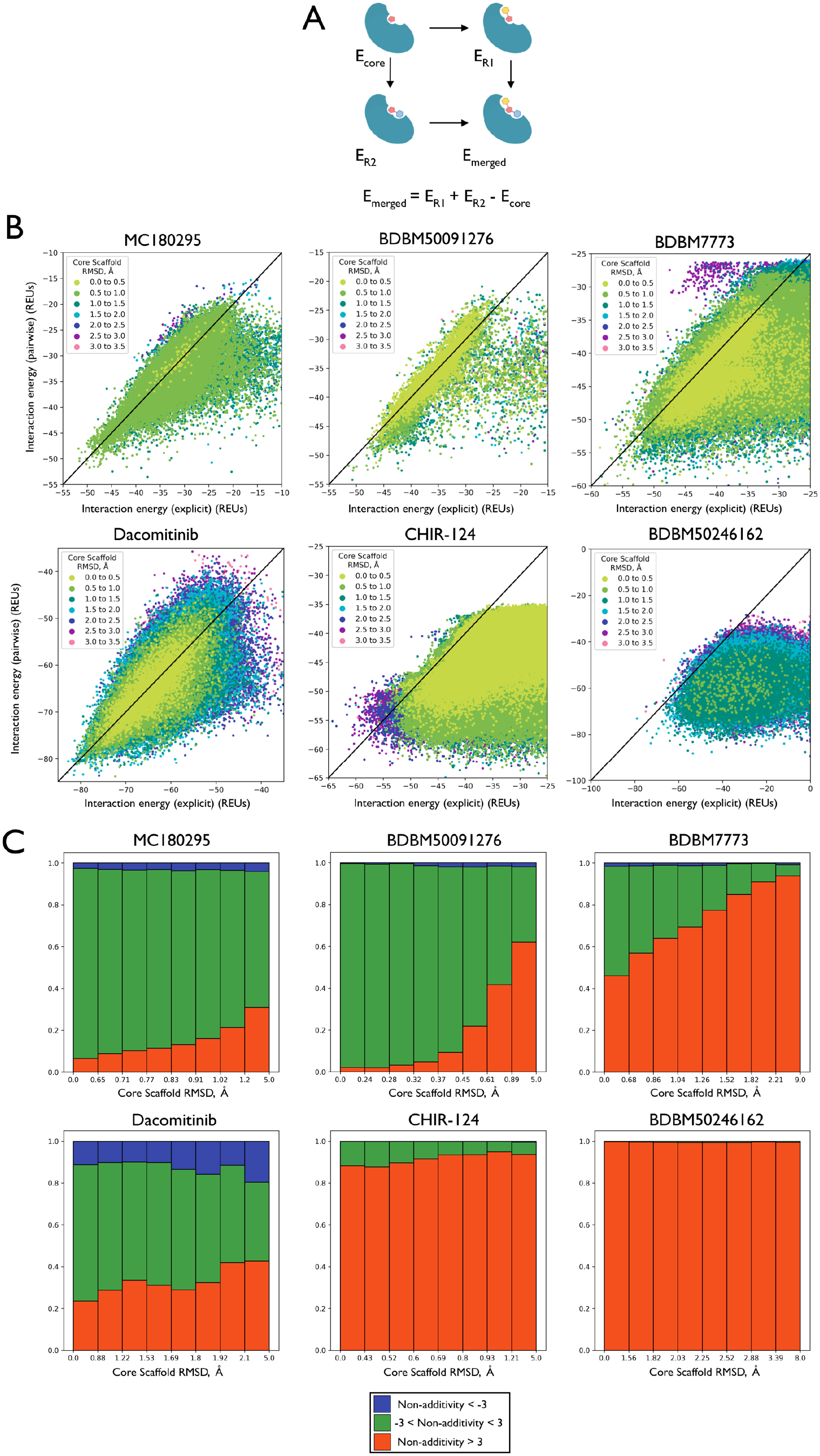
Pairwise additivity allows the inhibitors’ interaction energies to be predicted from their component fragments. (**A**) Chemical double-mutant cycle for estimating the merged inhibitors’ interaction energy. Subject to the pairwise additivity approximation, the interaction energy of the merged compound (E_merged_) is given by the sum of the fragments’ interaction energies (E_R1_, E_R2_) minus the interaction energy of the hinge-binding core (E_core_) which is present in both fragments. (**B**) For each compound in the corresponding enumerated library (**Table S4**), interaction energies are reported either by estimating from the component fragments using the pairwise approximation (y-axis), or by calculating explicitly from the re-refined model of the complete inhibitor (x-axis). Energies match closely (*points along the diagonal*) provided that the RMSD is small between the fragments’ positioning of their shared core (*light green points*). As the RMSD becomes larger (*blue and purple points*), the interaction energy of the complete inhibitor is often not as favorable as expected from the pairwise approximation (*points below the diagonal*). Interaction energies are reported in Rosetta energy units (REUs). (**C**) Within each of the six libraries, points are binned based on the component fragments’ RMSD with one another (bins are defined such that each bin contains 1/8 of the library). Within each bin, the pairwise-approximated interaction energy is subtracted from the explicitly-calculated interaction energy. The proportion of compounds for which the additivity approximation hold closely (within 3 Rosetta energy units) are shown in *green*. The proportion of compounds for which the explicitly-calculated interaction energy is worse than expected by the pairwise approximation are shown in *red*, and compounds better than expected are shown in *blue*.

Results from this experiment show that for our first four testcases, the interaction energy estimated with the pairwise approximation closely matches the full inhibitor’s interaction energy calculated explicitly (**Figure 5b**), as observed through points that lie along the diagonal of these plots. Agreement between the two values is highest when the position of the core scaffold matches very closely among the two fragments, as expected. For the BDBM7773 library, some of the compounds have with worse interaction energy than expected based on additivity (points below the diagonal); however, these the points have relatively high RMSD (dark green rather than light green). Put another way, merged compounds with worse interaction energies than expected from the pairwise approximation occur when the fragments do not position the shared core in precisely the same location.

To further visualize this relationship, we collected the points within each RMSD range and determined their adherence to the additivity approximation (**Figure 5c**). We find that compounds built from fragments in which the core scaffolds are well-aligned (low RMSD) are more likely to have pairwise interaction energies in close agreement with their explicitly calculated interaction energies (*green*); as the RMSD becomes larger, an increasing proportion of the compounds have explicitly calculated interaction energies that are worse than expected by assuming additivity (*red*).

The lack of additivity is most striking for the CHIR-124 and BDBM50246162 libraries; these are the two collections which diversify at three substituent positions rather than just two. Interestingly, the origins of non-additivity in these two cases are different from one another. In the case of CHIR-124, there are numerous compounds built from fragments that place the core scaffold in overlapping orientations (i.e., many light green points in **Figure 5b**), and yet their explicitly calculated energies are worse than expected based on additivity. Examination of these cases showed that steric clashes between the substituents were responsible for the observed non-additivity, stemming from the fact that the R1 and R3 vectors for substitution of this scaffold are close to one another (**Figure 3**). Conversely, in the case of BDBM50246162 low RMSDs are rarely sampled (i.e., there are very few light green points in **Figure 5b**). In this case, the complexity of the core (as well as the availability of multiple hydrogen bonding groups) leads to very few fragment triplets that place the core in precisely the same position.

In a virtual screening context, however, the cost associated with inaccurate scoring is not evenly distributed for the entirety of the library. If a compound is (correctly) deemed unworthy of advancement, then its score is irrelevant; provided these compounds receive poor scores, their precise values are irrelevant. Conversely, errors for the top-scoring compounds in the library can be much more costly, since this can greatly affect which compounds are advanced for further characterization.

Thus, the most relevant evaluation of the pairwise additive approximation is instead to determine whether the top-scoring compounds are indeed prioritized within the library, mimicking the primary objective of a virtual screen. For each library, we therefore selected the top 20 compounds on the basis for their explicitly calculated energies. We marked these top-scoring compounds as “active”, and the remainder of the library as “inactive”. We then evaluated the extent to which the “active” compounds were enriched in the top-ranking compounds selected using the pairwise additive scores.

Dramatically, we find that in all six cases the pairwise additive approximation allows early retrieval of the top-scoring compounds (**Figure 6**). Even in the two cases for which the complete library exhibited notable non-additivity (CHIR-124 and BDBM50246162), the top-scoring compounds can readily be identified using this approximation. In other words, the main effect of non-additivity in these two libraries applies to compounds that are not among the best in the library. The effectiveness at identifying the top-scoring compounds when using the pairwise additive approximation to screen these libraries – accompanied by typically about a 1000-fold reduction in computational demands – strongly supports the use of this approach for screening enumerated libraries of kinase inhibitors.

**Figure 6:**
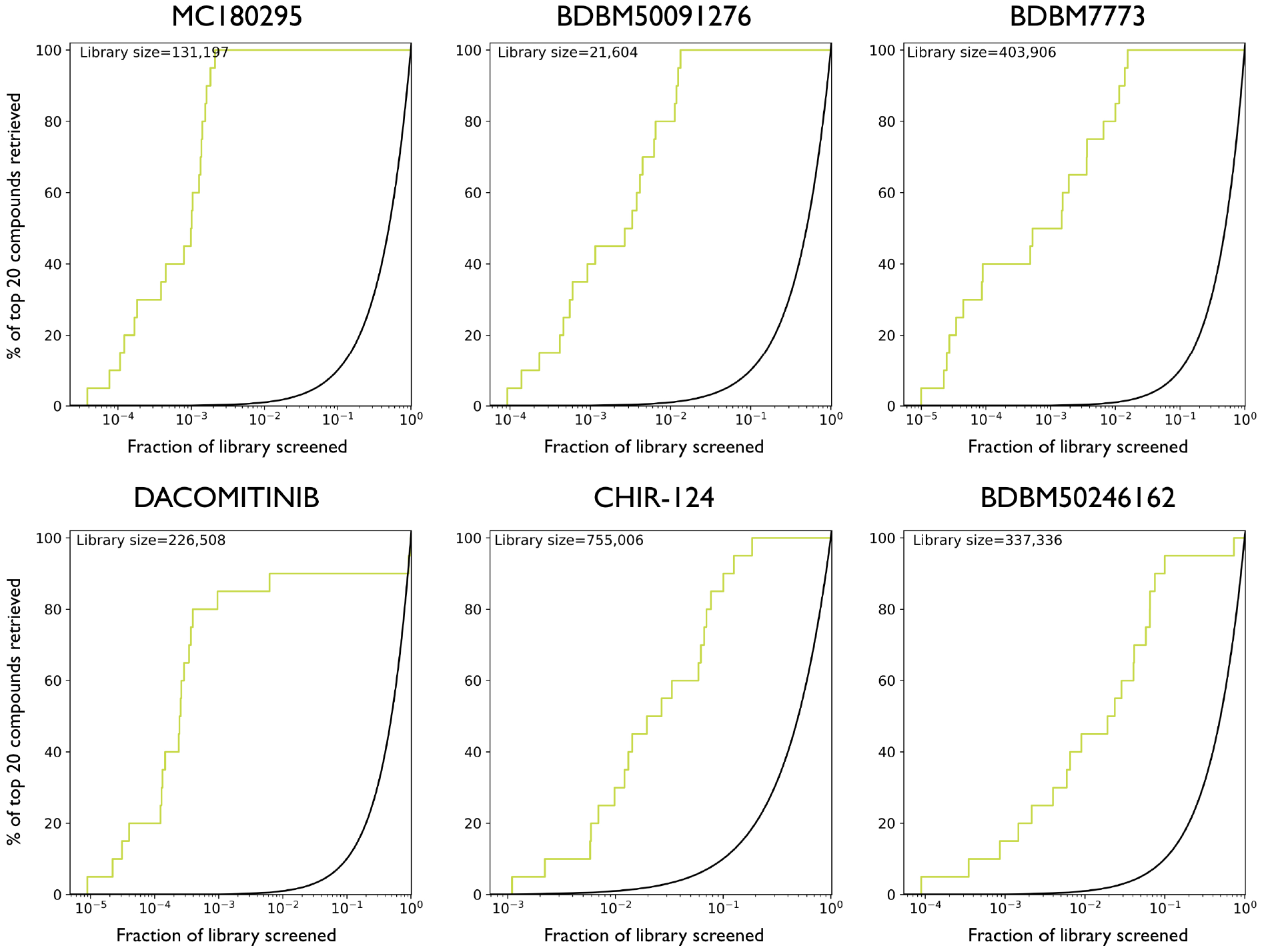
Rapid library-screening using the pairwise-additivity approximation, to find the top-scoring compounds. The 20 top-scoring compounds based on explicit screening were first identified from each library, and they were designated as “active” for the purpose of this benchmark experiment. The same library was then re-screened using the pairwise additive approximation. Receiver operating characteristic (ROC) curves show the percentage of “gold standard” compounds included in increasing subsets of the original library prioritized using the pairwise additive approximation. In all cases, the top-scoring compounds from the explicit screen were retrieved (*green curve*) much faster than expected by chance (*black curve*): in most cases, using the pairwise additive approximation allows more than half of the “gold standard” compounds to be identified within the top 1%. A very small number of compounds could not be minimized due to technical issues (clashes in the starting conformation), and these were excluded from consideration for this experiment (the size of the complete enumerated libraries are reported in **Figure S4**).

### Screening huge chemical libraries

It is important that the calculated interaction energies of kinase inhibitors can be reliably estimated from the interaction energies of its component fragments (provided that the cores are well-aligned with one another), because this implies that the original (huge) enumerated libraries need not be screened explicitly. Rather than build models for each compound in complex with the target kinase, one can instead build models only for the component *fragments*, and from these infer the interaction energies of the complete compounds. From a screening standpoint, one can thus identify the top-scoring compounds from these huge libraries extremely rapidly, and then use the prescribed synthetic route to make several compounds and test their activity.

While experimentally validating the top-scoring compounds lies beyond the scope of this first study, we note that the six inhibitors that inspired our study were themselves the culmination of campaigns that re-used the same synthetic route to explore many analogs. From the publications describing these inhibitors [26-28,31,32,56,57], we therefore selected the four most potent analogs reported in each study, and asked whether our method would prioritize these compounds from among the vast swath of enumerated chemical space.

We began by confirming that the fragments needed to build each of these analogs were present in our enumerated libraries. To avoid having to screen nearly a million building blocks in some cases, we then randomly excluded building blocks to reduce the size of each fragment collection (**Table 1**) while preserving those fragments needed for building the specific analogs reported in the literature. We then further filtered the fragment sets to remove large substituents (that would bring the final inhibitors molecular weight over 500 Da) and atom types not supported by the OMEGA software. For each library, we then built models corresponding to all fragments bound to the cognate kinase. Comparison of the fragments usage of the binding site sub-pockets (as defined by KinFragLib [55]) to that of the parent inhibitor, we confirmed that each fragment library provides alternate means of occupying essentially the same volume in the kinase active site as the parent inhibitor (**Figure S2**). We then evaluated the interaction energy for each fragment, and we used the pairwise additive approximation to rapidly estimate the interaction energy for all compounds in these huge libraries.

**Table 1:**
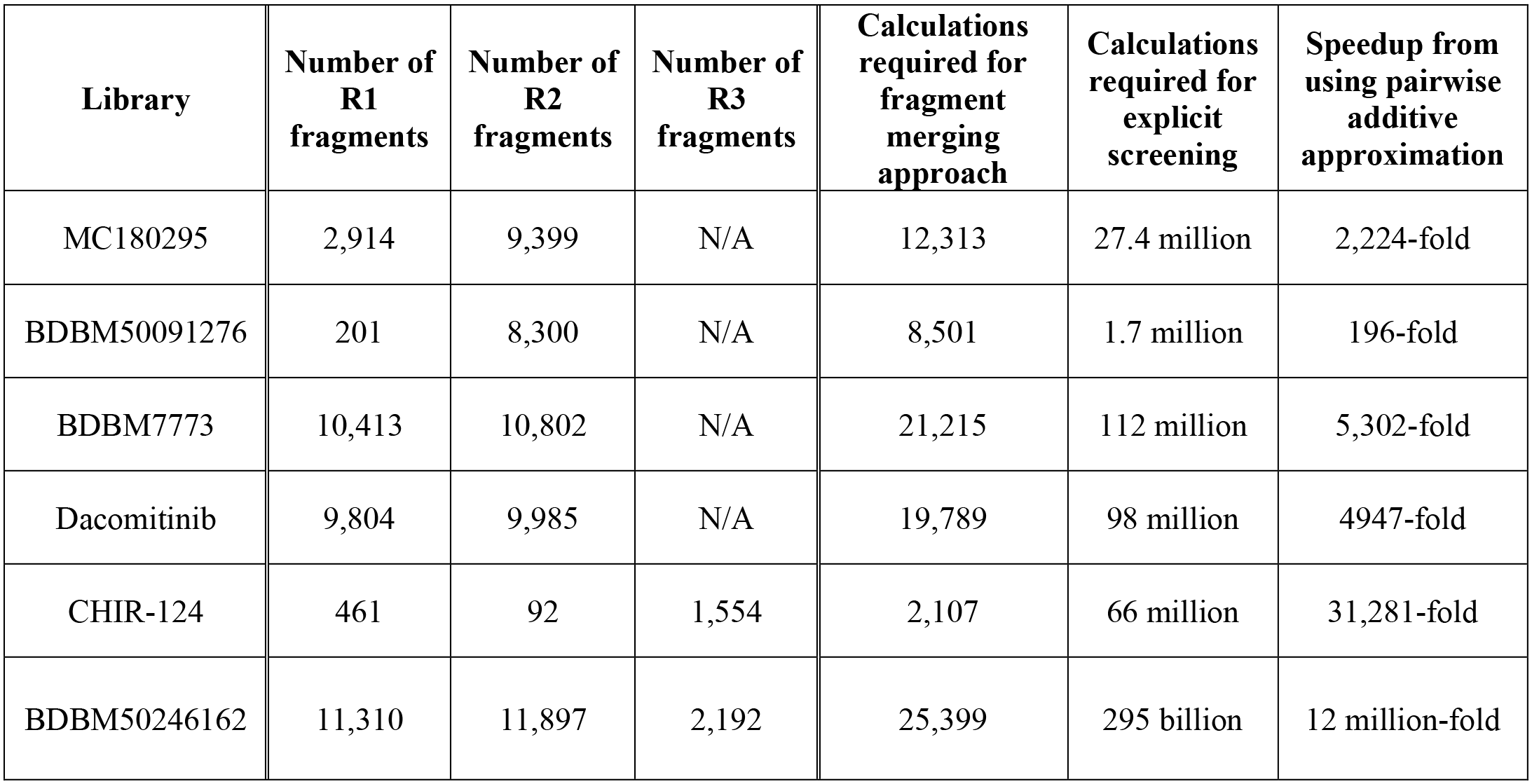
Large libraries for carrying out realistic virtual screens. For the retrospective screening experiment in which we determine whether known active compounds are identified from the enumerated libraries, computational requirements scale as the number of building blocks (rather than the number of enumerated compounds). This enables screening of very large libraries (with energy minimizations that include the kinase internal degrees of freedom), using modest computational requirements. Explicitly screening the same libraries would require many orders of magnitude more computational resources, as highlighted in the rightmost column.

| Library | Number of R1 fragments | Number of R2 fragments | Number of R3 fragments | Calculations required for fragment merging approach | Calculations required for explicit screening | Speedup from using pairwise additive approximation |
| --- | --- | --- | --- | --- | --- | --- |
| MC180295 | 2,914 | 9,399 | N/A | 12,313 | 27.4 million | 2,224-fold |
| BDBM50091276 | 201 | 8,300 | N/A | 8,501 | 1.7 million | 196-fold |
| BDBM7773 | 10,413 | 10,802 | N/A | 21,215 | 112 million | 5,302-fold |
| Dacomitinib | 9,804 | 9,985 | N/A | 19,789 | 98 million | 4947-fold |
| CHIR-124 | 461 | 92 | 1,554 | 2,107 | 66 million | 31,281-fold |
| BDBM50246162 | 11,310 | 11,897 | 2,192 | 25,399 | 295 billion | 12 million-fold |

Ultimately, this approach allowed us to rapidly screen libraries ranging from 1.7 million compounds to 305 billion compounds, while allowing for protein flexibility in response to the different inhibitors’ substituents. To reiterate the effect of the pairwise additive approximate on the computational demands of these screen, we use as a representative example our screen of the MC180295 library. For each fragment, up to 10 conformers are separately minimized in complex with the kinase (the degrees of freedom are restricted to the substituent, making a small number of conformers is sufficient). A total of 2914 fragments were considered for the first substituent (29012 conformers), and 9399 fragments (90085 conformers) for the second substituent. Each conformer was minimized in complex with CDK9, for a total of 119,097 independent minimizations. These took an average of 2.5 min on relatively slow CPU’s, for a total of 4962 CPU-hours. By running these calculations on a typical cluster, the complete set of fragment interaction energies can thus be collected in a matter of hours (elapsed real time, aka “wall time”). Using this modest calculation, we can then infer interaction energies for all 27.4 million compounds (2914 x 9399) that comprise our enumerated library. While fast docking methods have been used to screen even larger libraries than this [7], these have required vastly larger computational resources and have screened against a fixed protein conformation: by contrast, the minimization step used here allows the protein to adjust its conformation in response to different ligands.

The interaction energy calculated using Rosetta [48, 49] comprises a sum over pairs of atoms, and consequently it tends to strongly favor larger ligands over small ones (by virtue of having more atoms, large ligands simply accumulate more interactions). The same trend broadly holds for experimentally-measured binding affinities as well (larger compounds tend to have better potency), which is problematic because larger compounds may be less advanceable due to future problems with absorption, distribution, metabolism, or excretion (ADME); accordingly, many optimization campaigns instead prioritize compounds on the basis of “ligand efficiency” (binding free energy divided by the number of non-hydrogen atoms) or related metrics [58–61]. To align with these goals, we ranked compounds in our enumerated screening libraries on the basis of their substituents’ calculated ligand efficiency (Rosetta interaction energy divided by the number of non-hydrogen atoms).

We present the relative ranking of the known analogs relative to the rest of the enumerated libraries using receiver operating characteristic (ROC) curves. For each model system, this provides a means to present the fraction of the library that ranks ahead of the each of the analogs known to be active against the kinase of interest (additional compounds published alongside the original inhibitor). While this is a standard approach for presenting retrieval of known active compounds in virtual screening benchmarks, though, it is not without well-understood biases [62–66]. Specifically, this analysis assumes that all “decoy” compounds (the compounds in the library not designated as “actives”) are inactive; thus, a good method should prioritize the known actives ahead of all the decoy compounds. If the decoys are clearly absurd with regards to their complementary for the protein target, they can be easily discarded and any method will appear to have excellent performance; thus, the difficulty of a given benchmark depends on the extent to which the decoys are suitably reasonable for the protein target and/or property-matched to the actives. With respect to suitably matched decoys, however, it is important to note that the “decoys” are typically compounds that have not even been tested for the activity of interest. If the likelihood of a decoy compound having activity is low, then the assumption that they are truly inactive may hold. On the other hand, if the decoy compounds are similar enough to the active compounds, they may themselves also have activity; these could then be (correctly) prioritized by the screening method, but they would be (incorrectly) penalized for doing so (because all decoys are assumed to be inactive).

In our experiment, the enumerated libraries bear the same core scaffold as the known actives, and many substituents have size and chemistry that is likely to make these active as well. For this reason, many of the decoy compounds in this experiment are themselves likely active; even a method providing “perfect” performance would not distinguish the actives that happen to have been experimentally characterized versus the undiscovered actives in our library. Thus, the layout of this experiment is such that a very conservative perspective of performance will be conveyed.

As a starting point, we first carried out this screen using the structure of the kinase that had been solved in complex with one of the reported inhibitors (**Figure 7**, *blue*). For all six cases, the known analogs were retrieved far sooner than would be expected by random chance (**Figure 7**, *green*). For a case with 20 active analogs, would expect to require a subset comprising 5% of the original library to find one by random chance. By contrast, for all six testcases we identify at least one of the known actives within the top-scoring 0.2% of the library. Gratifyingly, in all six cases we find that most of the known active compounds adhere closely to the pairwise additive approximation (**Figure S3**), consistent with their prioritization in this retrospective screening experiment.

**Figure 7:**
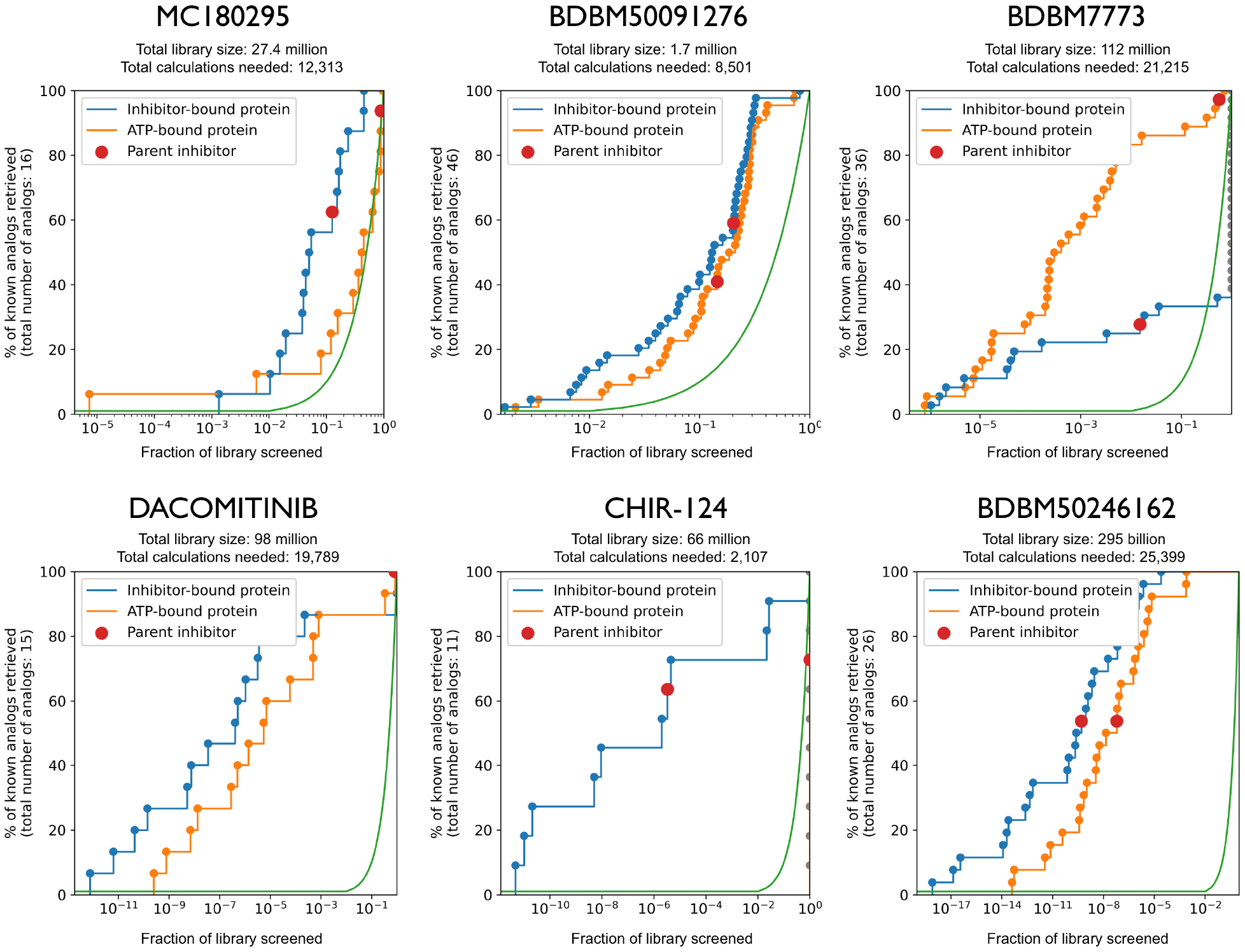
Large-scale screening of enumerated libraries using the pairwise additive approximation. For each of the six inhibitors in our set, we built large enumerated libraries (**Table 1**) and ranked the compounds on the basis of their substituents’ ligand efficiency (thus, scores arose in a pairwise additive way). Each library included six analogs reported to have activity against the target kinase. We plot the fraction of the library that must be screened before encountering each of these six active compounds (noting that many of the compounds ranked ahead of these six compounds may themselves also be active). In all six cases, the compounds reported in the literature to have activity are ranked more favorably within the library than expected by chance (*green curve*). Of the six analogs, one of them (*red dot*) corresponded to the inhibitor present in the crystal structure from which the kinase conformation was taken (*blue curve*). Screening was also separately carried out using a kinase conformation that was instead solved in complex with a nucleotide (various ATP analogs) (*orange curve*). The size of these libraries ranged from 1.7 million to 295 billion compounds (**Table 1**); by assuming pairwise additivity, these could be screened in a matter of hours on a typical cluster while including protein flexibility in response to the various substituents.

These first screens were carried out starting from a kinase crystal structure that was determined in complex with an inhibitor bearing the same hinge-binding motif as all compounds in the enumerated library. While structures of related compounds are likely to be available in a true hit-to-lead scenario, we nonetheless sought to examine performance when such a structure is not available. Accordingly, we carried out the same screens starting from nucleotide-bound crystal structures of the same kinases. We find that performance is only slightly diminished in this regime (**Figure 7**, *orange*), with one exception: for the CHIR-124 library, the fragments needed to build the reported analogs do not yield hinge-binding poses when using the ATP-bound protein structure, and they are thus not found in our screen.

We also sought to understand whether starting from a crystal structure solved in complex with one particular inhibitor would bias the screen to “rediscover” the compound used in solving the structure: if so, this would inadvertently focus hit-to-lead exploration of chemical space near the known structure. To test this, we evaluated the order in which the known analogs were ranked: in all but one case, the compound used for crystallography was *not* the top-scoring amongst the known analogs (**Figure 7**, *red dot*), confirming that our modeling protocol allows sufficient flexibility for the kinase structure to accommodate (and prioritize) alternate substituents.

The chemical structures of the analogs themselves further highlight the ability of this approach to recognize active compounds with diverse substituents (**Tables 2-7**). However, the redundancy that is evident among the analogs for a given target (the same R1 substituent is used with multiple R2’s, and vice versa) also serves as a reminder that many more combinations of these substituents are also likely to be active, if they can be combined using a shared positioning of the hinge-binding core. Because these compounds would not be designated as “active” in our ROC plots, the true performance of this approach is likely superior to that suggested by this retrospective analysis. Simply put, the chemical space available by enumerating these reactions is far larger than can be explored via “wet” medicinal chemistry.

**Table 2:**
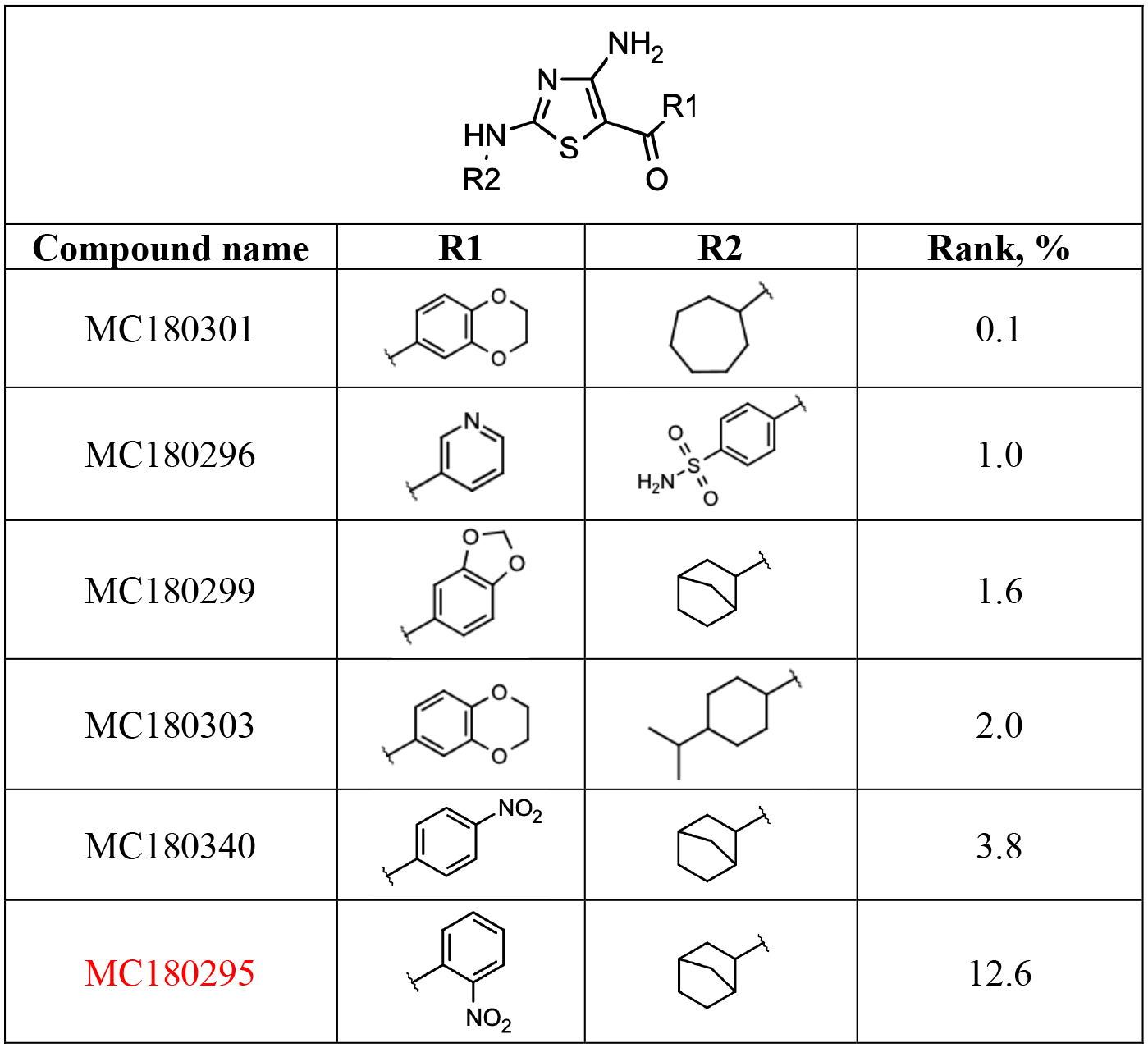
Among analogs of MC180295 previously reported to have activity against CDK9 [26], the top-scoring 5 as recovered from in our complete enumerated library are presented along with their ranking in this library (**Figure 7**). The compound present in the crystal structure (i.e., MC180295 itself) is indicated in *red*. The complete set of active analogs included in this experiment are included as Table S5.

**Table 3:**
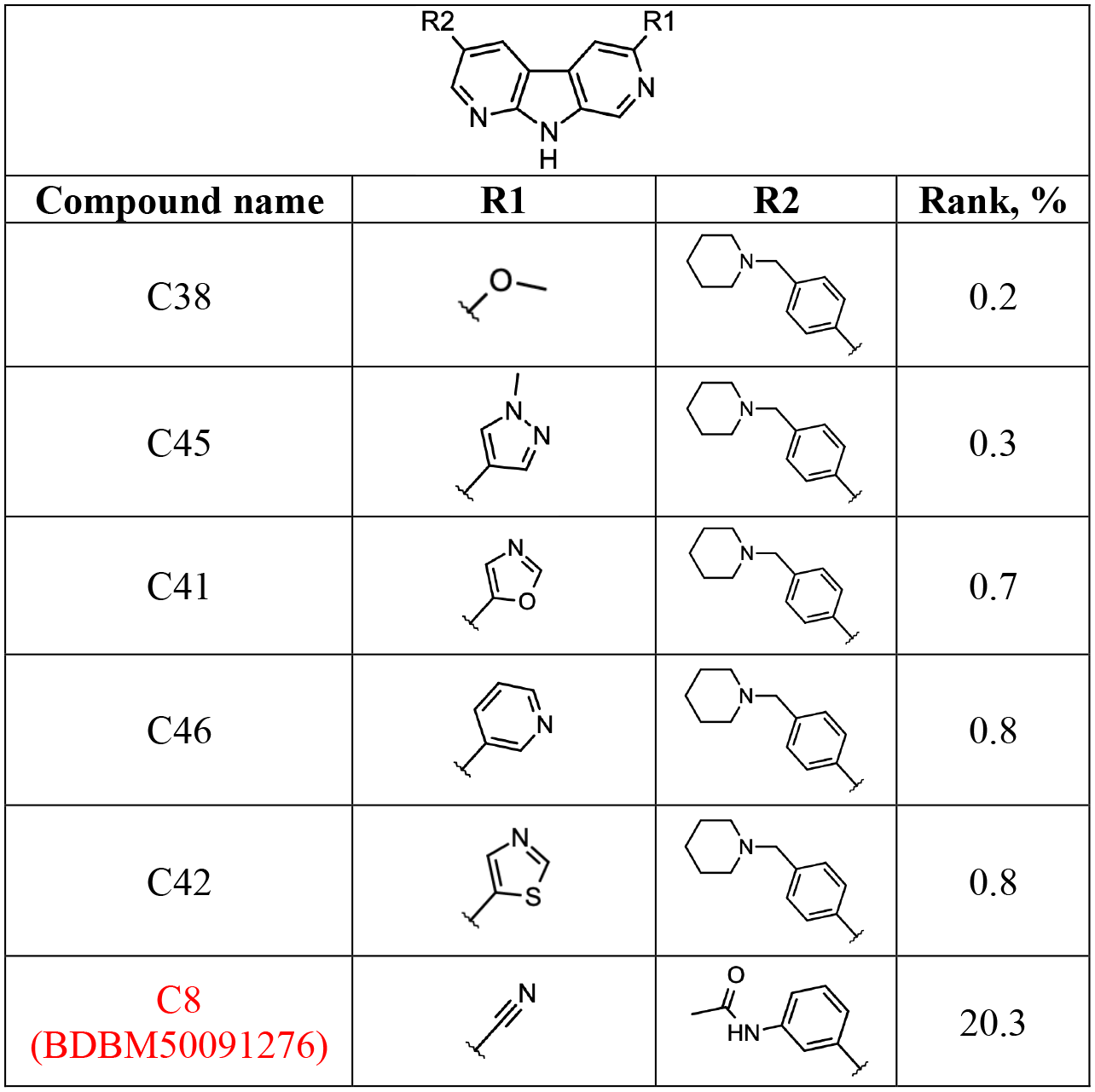
Analogs of BDBM50091276 previously reported to have activity against CHK1 [27]. The rank of each analog in our screen is provided relative to the complete enumerated library (**Figure 7**). The compound present in the crystal structure (i.e., BDBM50091276 itself) is indicated in *red*. The complete set of active analogs included in this experiment are included as Table S6.

|  |  |  |  |
| --- | --- | --- | --- |
| Compound name | R1 | R2 | Rank, % |
| C38 |  |  | 0.2 |
| C45 |  |  | 0.3 |
| C41 |  |  | 0.7 |
| C46 |  |  | 0.8 |
| C42 |  |  | 0.8 |
| **C8**<br>**(BDBM50091276)** |  |  | 20.3 |

**Table 4:**
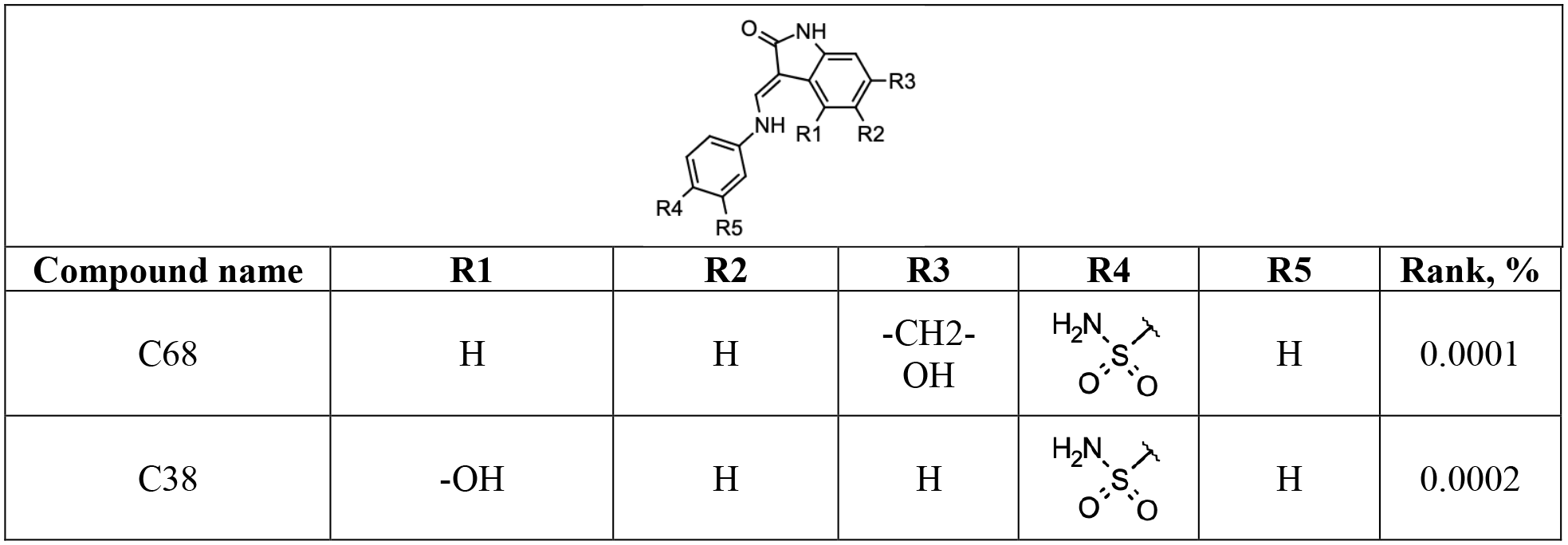
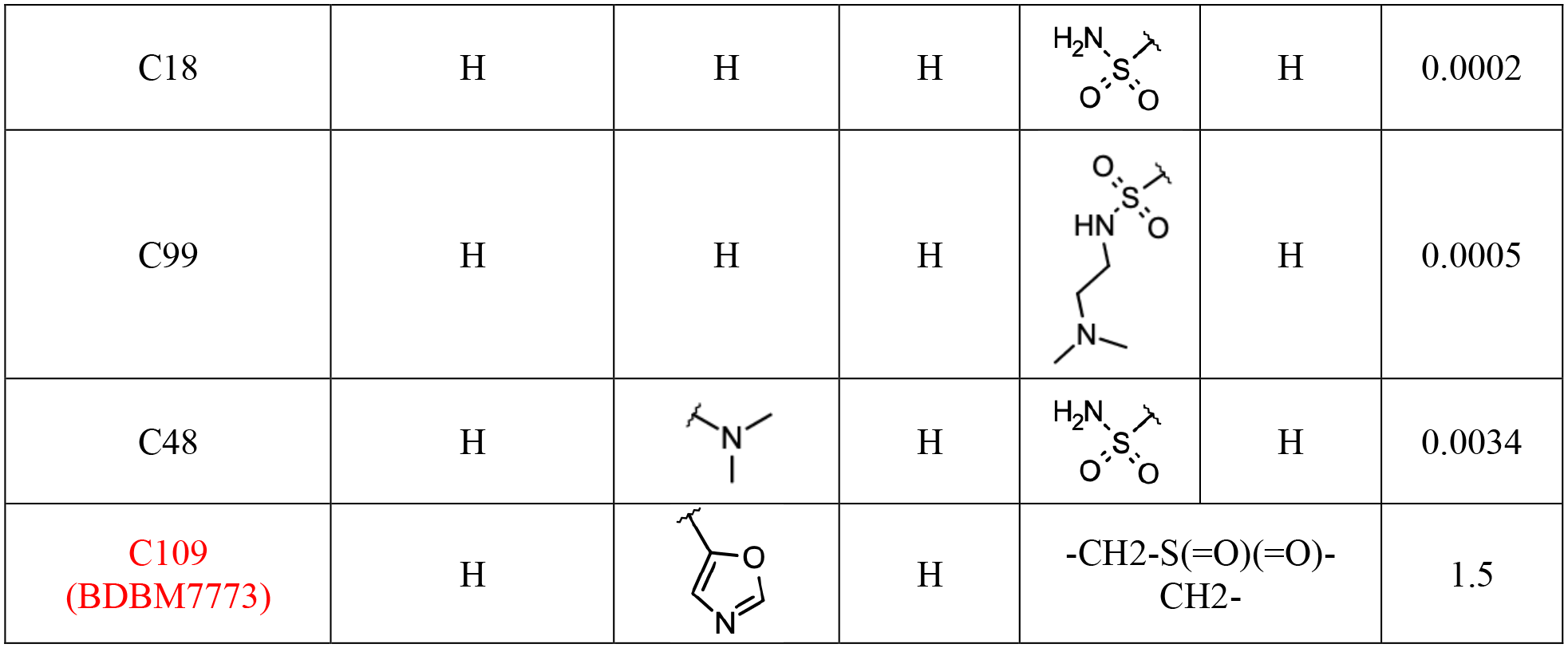
Analogs of BDBM7773 previously reported to have activity against CDK2 [28]. The rank of each analog in our screen is provided relative to the complete enumerated library (**Figure 7**). The compound present in the crystal structure (i.e., BDBM7773 itself) is indicated in *red*. The complete set of active analogs included in this experiment are included as Table S7.

**Table 5:**
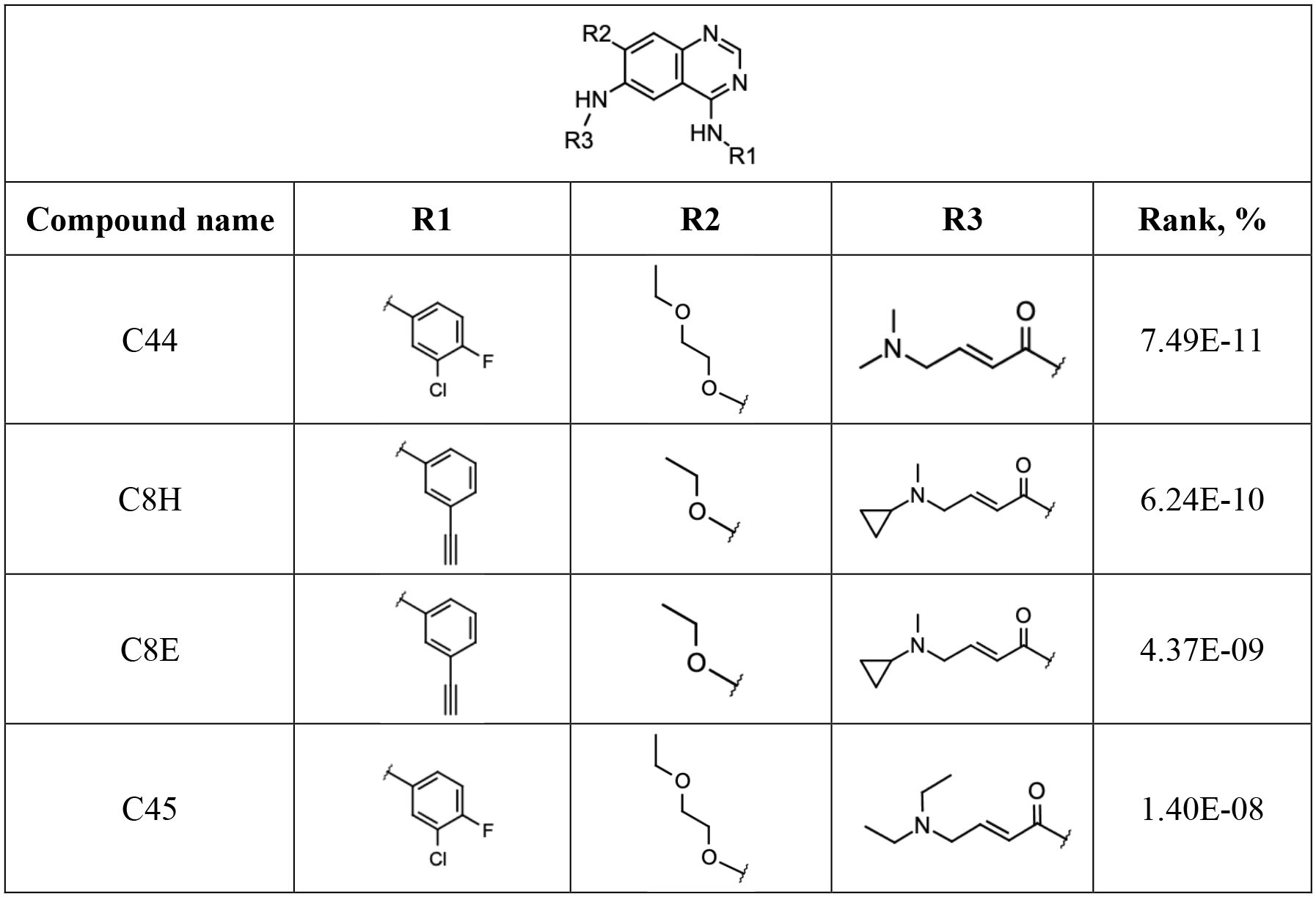
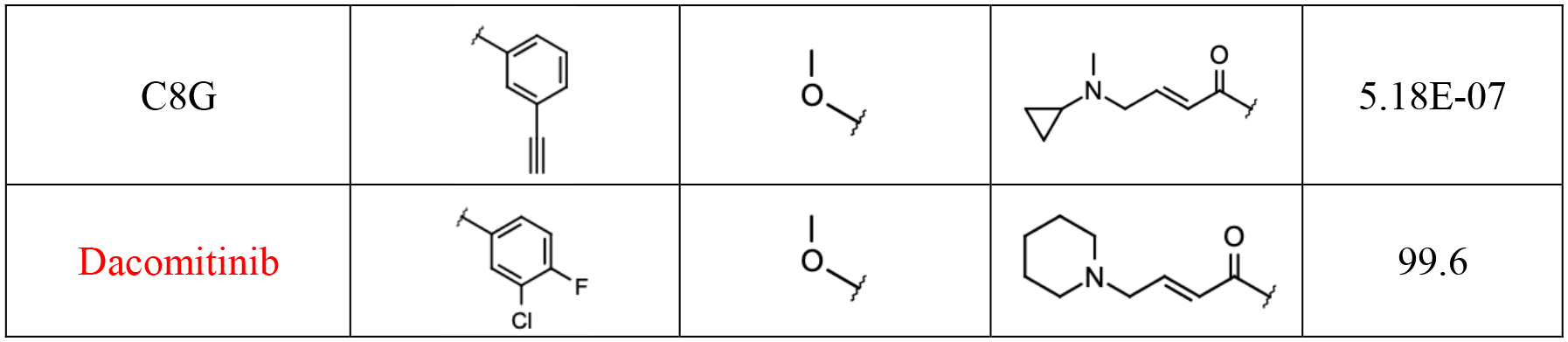
Analogs of Dacomitinib previously reported to have activity against EGFR^T790M^ (or EGFR itself) [56, 57]. The rank of each analog in our screen is provided relative to the complete enumerated library (**Figure 7**). The compound present in the crystal structure (i.e., Dacomitinib itself) is indicated in *red*. The complete set of active analogs included in this experiment are included as Table S8.

**Table 6:**
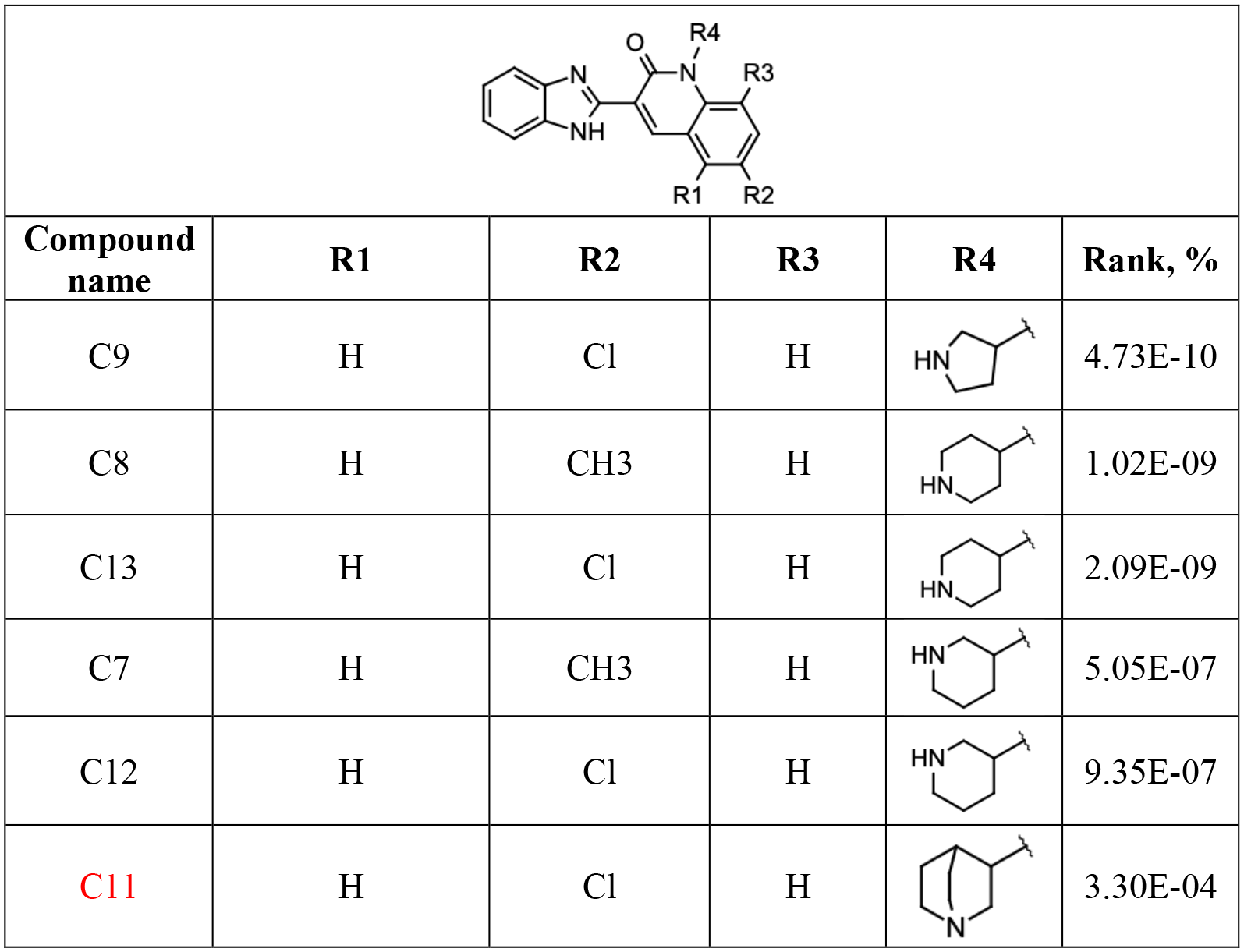
Analogs of CHIR-124 previously reported to have activity against CHK1 [31]. The rank of each analog in our screen is provided relative to the complete enumerated library (**Figure 7**). The compound present in the crystal structure (i.e., CHIR-124 itself) is indicated in *red*. The complete set of active analogs included in this experiment are included as Table S9.

**Table 7:**
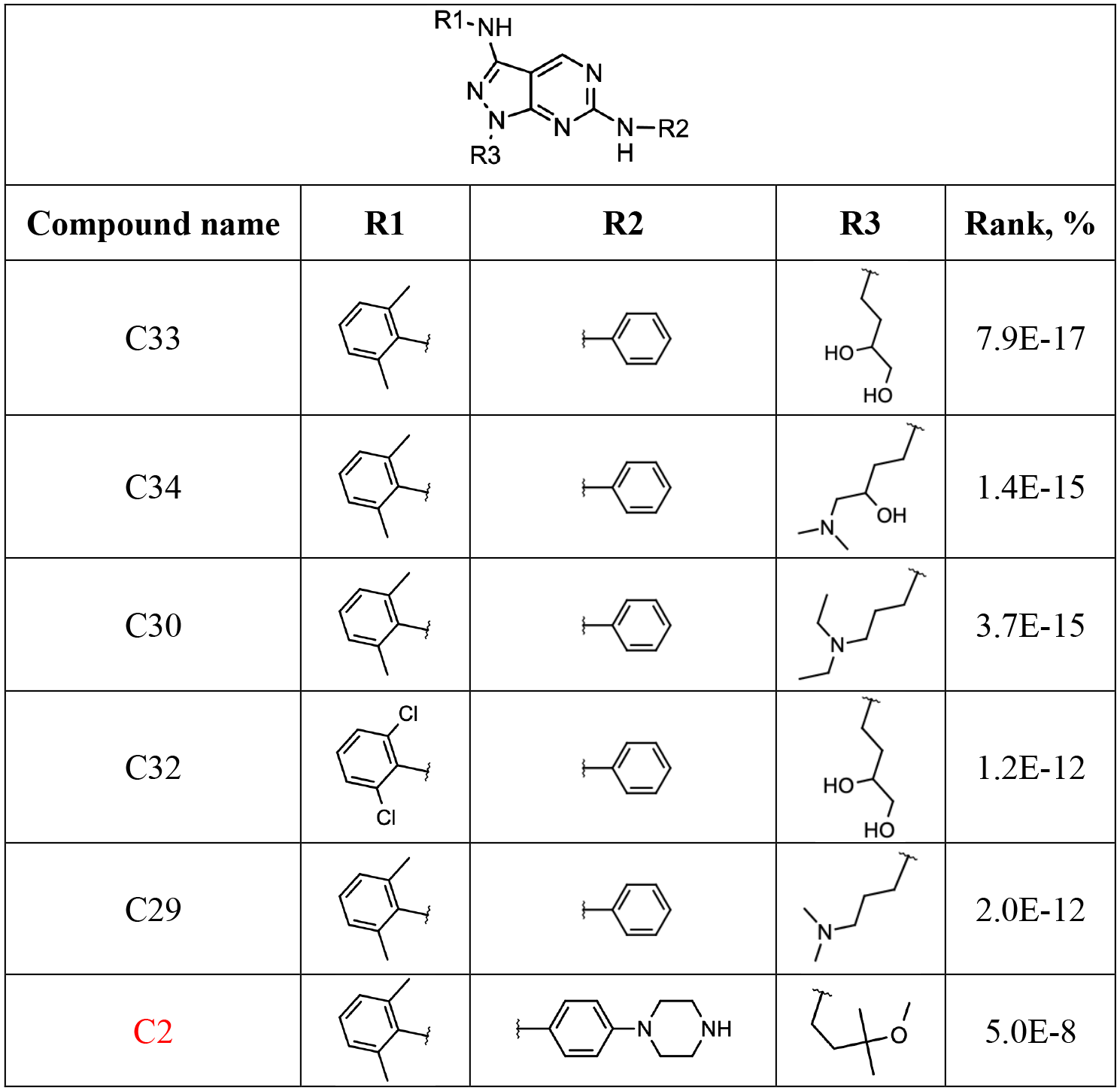
Analogs of BDBM50246162 previously reported to have activity against ACK1 [32]. The rank of each analog in our screen is provided relative to the complete enumerated library (**Figure 7**). The compound present in the crystal structure (i.e., BDBM50246162 itself) is indicated in *red*. The complete set of active analogs included in this experiment are included as Table S10.

## Discussion

In light of the intense focus on kinases as drug targets, there has been extensive interest in streamlining hit-to-lead optimization for this target class – including through computational methods.

With respect to library building, for example, methods have been recently described for deconstructing sets of reported kinase inhibitors, then reassembling these into new collections for screening. One such method operates at the level of chemical structure, by assigning each piece of a known inhibitor to be either a “core” (hinge-binding) fragment, a “connecting” fragment, or a “modifying” fragment. These fragments were then recombined to yield a library of 39 million compounds, albeit with the caveat that the three-dimensional fit of these compounds with respect to the ATP-binding site has not been established [67]. Addressing this, a complementary method instead elected to divide the ATP-binding site into six sub-pockets, and then collect from the PDB each of the various fragments that have been observed to occupy these regions. The authors then recombine fragments drawn from different structures, to yield a library of nearly 7 million new compounds [68]. In both cases, the resulting libraries are highly enriched in novel chemical matter, but synthetic accessibility of the generated compounds is not assured. By contrast, the libraries that we present here are vastly bigger than these earlier libraries, because we instead enumerate compounds that can be constructed using commercially-available building blocks, rather than simply using pieces of existing kinase inhibitors. While not guaranteed, the likelihood that the compounds in our enumerated libraries will be synthetically facile is also much higher, because each compound is included specifically because it is likely to be amenable to assembly using a pre-determined synthetic route.

Beyond just kinase-focused methods, more general strategies for de novo design of new compounds have been broadly explored for more than 30 years. Whereas early methods proposed for this task often produced compounds for which synthetic tractability was a major limitation [69], a number of more recent approaches have been validated in retrospective and prospective evaluations to confirm both synthetic tractability and activity for the biological target of interest [70]. New methods for growing ligands into a (fixed) protein environment continue to be an active focus of research, particularly using fragment replacement strategies [71], template-driven docking [72], genetic algorithms [73], and deep learning [74]. Modern methods can yield impressive performance, though typically with the caveat that benchmarking typically entails building back a ligand into the protein structure it was taken from – so, it remains unclear if (or how well) these methods can accommodate changes to the protein structure in response to the ligand.

Overall, our library-building strategy is most similar to that encoded as part of the Diversity-Oriented Target-focused Synthesis (DOTS) workflow [75]. This iterative hit-to-lead strategy starts from the crystal structure of an initial fragment hit, and it considers chemical transformations (using available building blocks) that could be used to elaborate the fragment. The elaborated fragments are ranked using virtual screening, then the top-scoring compounds are synthesized and tested (using robotic instrumentation for the synthesis). The top-performing derivatives are then used as the starting point for a new round of optimization. Another related approach, PathFinder [76, 77], uses free energy perturbation (FEP) calculations to evaluate synthetically-tractable substituents, coupled with active learning to reduce the computational requirements by inferring effects of substitutions that have not yet been explicitly modeled.

Using enumerated libraries, it is easy to very quickly build up chemical libraries that become too large for explicit virtual screening. This has already been observed in the context of computational hit finding (i.e., naïve screening, as opposed to hit-to-lead optimization), enabled by the growth of “make-on-demand” catalogs from vendors including Enamine [7]. Back when this library comprised “only” 170 million compounds, it was possible to explicitly dock each compound through a massive brute-force campaign [7]. By now, the library has grown to 19 billion compounds available for delivery, which far exceeds what can be addressed through any kind explicit structure-based screening. Our own previous work explicitly screened diverse representatives of an intractably large library to search for useful chemotypes first, then extracted from the library analogs of these compounds for a second round of screening [13]. Other studies have trained machine learning models to predict docking scores from chemical structures, as a means to rapidly obtain the outcomes from docking in a much less resource-intensive way [78, 79].

The key finding of our study is that we can quickly and reliably estimate the interaction energy for the compounds in our enumerated libraries, by assuming pairwise additivity of the substituents. This finding is important, because it allows us to assign energies across huge libraries, without needing to explicitly dock each compound: only the component fragments need to be docked, and the compatibility of their poses verified. With respect to hit-to-lead optimization, it also confirms that one can find the compounds with the best interaction energies by simply selecting the best fragments and merging them with one another – but only if the fragments’ shared cores are precisely aligned with one another in the binding site.

Traditional hit-to-lead optimization in medicinal chemistry focuses on building up structure-activity relationships by evaluating the functional consequence of separately changing individual side chains: the underlying hope is that after multiple substituents are separately optimized, they can later be productively merged. This expectation is quantitatively expressed through Free Wilson analysis, which uses observed structure-activity relationships (SAR) to assign a specific contribution to each substituent [80]. Unfortunately, this conceptually pleasing method for deconvolution can frequently turn out to be insufficient, as substituents’ contributions can are often be non-additive [81]. At first, this seems to run counter to the claim we make here, that fragment contributions *are* strongly additive.

A recent study carried out a careful survey of examples from the literature that demonstrate strong and verifiable non-additivity, in a variety of different protein-ligand systems for which crystal structures are available [52]. In nearly every example identified, the authors find that the ligand exhibits an altered binding mode in response to one or more of the substitutions. Thus, the conclusion from these examples aligns closely with our own observation, that additivity can be assumed only if the shared core does not change position. While it is not feasible to use x-ray crystallography to verify the binding modes of all fragments in a hit-to-lead optimization campaign, an inherent advantage of using structure-based modeling for driving SAR is that the binding modes are provided for each fragment. This, in turn, is necessary to inform on which fragments can be productively merged with one another.

In our screens, a relatively high proportion of the fragment cores could be merged with one another. In part, this was due to our decision to focus at the outset on kinase inhibitors: as noted earlier, these inhibitors engage the kinase hinge region in a set of prescribed hydrogen bonds (**Figure 1b**). This interaction is very strongly templated, and thus tends to be closely recapitulated amongst the fragments that elaborate the hinge-binding scaffold. Thus, the architecture of the kinase binding site contributes to the conserved pose of the shared core, which in turn leads to additivity of the substituents. Conversely, and in keeping with the survey of examples in which non-additivity is observed [52], we do not expect that the same high proportion of fragments can be merged with one another for binding sites that lack a strongly templated interaction to enforce the pose of the shared core. In such cases, we expect that additional filters will be necessary to gauge which fragment pairings may be suitably merged with one another: beyond simply confirming the shared alignment of the common core, we envision that filtering to ensure the merged inhibitor can be built without extreme strain or intramolecular clashes will also prove useful [82].

While fragment pose prediction has historically proved challenging, improved methodologies have now delivered dramatic successes [83–85]. Looking forward, we expect that continued advances will soon allow for reliable identification of fragments that share overlapping cores. Such fragments can naturally be merged on the basis of their shared core, and then libraries to diversify the component fragments will provide a natural path for hit-to-lead optimization.

Finally, we leave with the caveat that the pairwise additive approximation allows for very rapid screening of huge chemical libraries on the basis of interaction energies. The use of interaction energies fundamentally ignores contributions from binding conformational entropy, and calculations are carried out in the absence of explicit solvent: accordingly, one cannot expect these interaction energies to strictly correlated with the compounds’ binding affinities. Much akin to deep learning models for predicting docking scores [78, 79], this strategy merely provides a way to rapidly estimate an admittedly-crude quantity (interaction energy) that cannot be evaluated at the scale needed to keep up with the growth of library sizes. It will prove very natural to integrate this strategy into a layered approach, by which the size of the library is progressively culled through the use of increasingly-expensive methods, and ultimately using tools such as free energy perturbation (FEP) calculations [86] and orthogonal machine learning scoring functions [13], in combination with tools for predicting ADME-PK [87], to select compounds for further study.

## Data and Software Availability

Computational tools to enable the use of this protocol are publicly available on GitHub (https://github.com/karanicolaslab/CombiChem).

Results described in our study relied on OMEGA, QUACPAC, RDKit, and PyRosetta as external dependencies. OMEGA version v3.0.0.1 and QUACPAC version 2.0.2.2 were used for these studies (OpenEye offers a free public domain license for purely non-commercial research, which can be used to obtain OMEGA and QUACPAC). RDKit version 2020.03.3.0 was used for these studies (RDKit is freely distributed under a BSD 3 license). PyRosetta version 2020.03.post.dev+57.master was used for these studies (PyRosetta is licenses are free for academic use).

## Methods

### Computational protocol

For a given fragment, protonation states and tautomeric states were assigned using OpenEye QUACPAC FixpKa, using pH 7.4. Partial charges assigned were in OpenEye’s QUACPAC (assigncharges.py, using AM1BCC). For each fragment, the charges for the shared core were taken from the charges assigned to the parent inhibitor: when merging fragments, we found that consistency between the partial charges was necessary for pairwise additivity. Because of this step, the formal charge on a given fragment (or merged inhibitor) was not necessarily an integer value.

Conformers were generated using OpenEye OMEGA. For our analysis of pairwise additivity, we generated up to 100 conformers for each fragment, and used an RMS threshold of 0.6 Å for conformer deduplication. For our screens, we instead generated 10 conformers for each fragment, and used the flags “-fixfile” and “-deleteFixHydrogens false” to keep the hinge-binding core fixed (thus restricting conformational diversity onto the substituent).

Alignments into the protein structure were carried out using RDKit, based on the core hinge-binding motif. Structures were then refined using PyRosetta’s MinMover for energy minimization, with the L-BFGS gradient minimizer at a convergence threshold of 1E-0.6, and the interaction energies were reported by PyRosetta.

Structures of the kinase-bound fragments were merged into the complete inhibitor using a Python script (included in the GitHub repository above). Finally, the complete inhibitor (after merging) was re-minimized using the same PyRosetta protocol, and the interaction energies were collected.

Modeling the dacomitinib-inspired library required extra consideration, because of the covalent linkage to the kinase. Rather than simply keep the conformation of the hinge-binding core fixed in conformer generation, the fixed region also included the complete side group harboring the Michael acceptor (**Figure 3**, region shown in *red*). Thus, each of the generated conformers could be appropriately linked back to Cys797, provided that the hinge-binding motif was correctly placed. Initial alignments for the resulting fragments proceeded as with each of the non-covalent testcases. A Rosetta parameter file was generated for the Cys adduct corresponding to each fragment, to allow minimization of the adduct (rather than the free fragment). Minimizations were then carried out as with the non-covalent testcases.

### Steps to ensure pairwise additivity of fragment scores

Our initial attempts to use the chemical double mutant cycle presented earlier (**Figure 5a**) did not yield strongly pairwise additive interaction energies. Careful decomposition of the contributions to the evaluated energies pointed to four factors that were responsible for essentially all of the non-additivity (**Figure S4**).

To obtain interaction energies that are closely pairwise additive, we found the following steps to be necessary:

1) <u>Consistent parametrization:</u> Partial charges were often assigned differently between the full inhibitor and the corresponding fragments. For this reason, we generated partial charges only using the full inhibitor, then transferred these charges to fragments.
2) <u>Avoiding steric clashes between substituents:</u> Scenarios can arise in which two different fragments present a substituent that occupies the same region of the binding site. When merged, the two substituents obvious clash with one another. For the examples presented in our study, this scenario occurred only in a very small number of cases (involving long and flexible substituents), because the substituted vectors on these hinge-binding cores point away from one another.
3) <u>Positioning of the hinge-binding core:</u> If two substituents each prefer a conformation of the shared core that is inconsistent with the other substituent, then both cannot be satisfied in the context of the merged compound. As presented earlier (**Figure 5b**), this source of non-additivity can be greatly mitigated by merging only pairs of fragments that position the hinge-binding motif in precisely the same pose.
4) <u>Protein conformation:</u> Similarly, if two substituents each require that the protein adopts a conformation that is inconsistent with the other substituent, then both cannot be satisfied in the context of the merged compound. As noted earlier, kinase inhibitors typically have a flat hinge-binding core, with substituents that extend in both directions parallel to the hinge (**Figure 1**). The substituent facing the kinase’s αC-helix occupies the so-called “hydrophobic back-pocket” of the binding site [88] (**Figure 1b**, *right side of the compounds drawn*), and can be associated with structural changes in response to the different inhibitors. By contrast, substituents at the other side of the hinge-binding core are typically partially exposed. Thus, modeling fragments that face the αC-helix often lead to small structural changes, whereas those facing in the opposite direction do not. When calculating energies for a complete inhibitor, we accordingly find that pairwise additivity holds more reliably when refinement is started from a protein structure that was pre-built in complex with the fragment that occupies the back-pocket.

### Estimating rank of compounds from fragment scores

In the benchmark screening experiments, we ranked experimentally-validated analogs relative to the rest of the enumerated library. While this was straightforward with smaller libraries, the library built around BDBM50246162 contained almost 300 billion compounds. Determining the precise rank for each active compound would require explicitly counting how many compounds from the library score better/worse than the active analog (a large calculation).

Instead, we instead estimated the rank of each active analog by relying again on the pairwise additivity of the components. For a given analog, we determined the rank of each of the three component fragments relative to all substituents at the corresponding position. For a given fragment, we converted the rank to a percentile (by dividing by the number of fragments), and then to a Z-score by using a standard normal table. Given the independence of the three component fragments, we then summed the Z-scores to yield a Z-score for the complete compound. We then used a standard normal table to convert the latter Z-score to a percentile, which provides an estimate of the proportion of the library that scores ahead of the query (active) compound.

### PDB structures used in calculations

Calculations involving structures of the parent inhibitors bound to their cognate kinases were drawn from the corresponding PDB structures: MC180295/CDK9 (PDB: 6W9E), BDBM50091276/CHK1 (PDB: 4RVK), BDBM7773/CDK2 (PDB: 1KE7), Dacomitinib/EGFR^T790M^ (PDB: 4I24), CHIR-124/CHK1 (PDB: 2GDO), and BDBM50246162/ACK1 (PDB: 3EQR).

Selectivity calculations across the CDK kinase family were initiated from crystal structures these proteins: CDK1 (PDB: 5LQF), CDK2 (PDB: 4EOQ), CDK5 (PDB: 4AU8), CDK6 (PDB: 2EUF), CDK7 (PDB: 1UA2), CDK8 (PDB: 5IDN), CDK9 (PDB: 3BLQ). Because there were no crystal structure available for CDK3, CDK4, and CDK11, comparative models were generated using SWISS-MODEL [89] for these (using PDB templates 4EOR, 1XO2, and 6GUE, respectively).

Screens were carried out against the inhibitor-bound kinases, and also against structures solved with a nucleotide (ATP/ADP/AMP/etc.) rather than an inhibitor. The nucleotide-bound structures used in these screens were: MC180295/CDK9 (PDB: 4IMY), BDBM50091276/CHK1 (PDB: 7AKM), CHIR-124/CHK1 (PDB: 7AKM), BDBM7773/CDK2 (PDB: 2CCH), and BDBM50246162/ACK1 (PDB: 1U54).

## Associated Content

Supporting Information. **Supporting Figure S1**, analysis of how fragments and merged compounds occupy the binding site. **Supporting Figure S2**, analysis of how large fragment libraries occupy the binding site. **Supporting Figure S3**, analysis of pairwise additivity for known active analogs. **Supporting Figure S4**, summary of reasons for lack of pairwise additivity. **Supporting Table S1**, SMARTS templates for identifying building blocks and transforming them into fragment libraries. **Supporting Table S2**, Search parameters for identifying building blocks in PubChem, using SMARTS matching. **Supporting Table S3**, Vendors included in our search for available building blocks. **Supporting Table S4**, the smaller libraries amenable to explicit screening, used in the study to evaluate additivity. **Supporting Table S5**, Complete set of known active analogs for MC180295 library. **Supporting Table S6**, Complete set of known active analogs for BDBM50091276 library. **Supporting Table S7**, Complete set of known active analogs for BDBM7773 library. **Supporting Table S8**, Complete set of known active analogs for Dacomitinib library. **Supporting Table S9**, Complete set of known active analogs for CHIR-124 library. **Supporting Table S10**, Complete set of known active analogs for BDBM50246162 library. This material is available free of charge via the Internet at http://pubs.acs.org.

## Acknowledgements

This work used the Extreme Science and Engineering Discovery Environment (XSEDE) allocation MCB130049, which is supported by National Science Foundation grant number ACI-1548562. This work was supported by grants from the National Institute of General Medical Sciences (R01GM123336) and from the National Science Foundation (CHE-1836950). This research was funded in part through the NIH/NCI Cancer Center Support Grant P30CA006927.

## References

1. Kontoyianni M. Docking and Virtual Screening in Drug Discovery. Methods Mol Biol. 2017; 1647:255–66.

2. Irwin JJ, Shoichet BK. ZINC--a free database of commercially available compounds for virtual screening. J Chem Inf Model. 2005; 45:177–82.

3. Irwin JJ, Sterling T, Mysinger MM, Bolstad ES, Coleman RG. ZINC: a free tool to discover chemistry for biology. J Chem Inf Model. 2012; 52:1757–68.

4. Jorgensen WL. Efficient drug lead discovery and optimization. Acc Chem Res. 2009; 42:724–33.

5. Cournia Z, Allen B, Sherman W. Relative Binding Free Energy Calculations in Drug Discovery: Recent Advances and Practical Considerations. J Chem Inf Model. 2017; 57:2911–37.

6. Wang L, Wu Y, Deng Y, Kim B, Pierce L, Krilov G, Lupyan D, Robinson S, Dahlgren MK, Greenwood J, Romero DL, Masse C, Knight JL, Steinbrecher T, Beuming T, Damm W, Harder E, Sherman W, Brewer M, Wester R, Murcko M, Frye L, Farid R, Lin T, Mobley DL, Jorgensen WL, Berne BJ, Friesner RA, Abel R. Accurate and reliable prediction of relative ligand binding potency in prospective drug discovery by way of a modern free-energy calculation protocol and force field. J Am Chem Soc. 2015; 137:2695–703.

7. Lyu J, Wang S, Balius TE, Singh I, Levit A, Moroz YS, O’Meara MJ, Che T, Algaa E, Tolmachova K, Tolmachev AA, Shoichet BK, Roth BL, Irwin JJ. Ultra-large library docking for discovering new chemotypes. Nature. 2019; 566:224–9.

8. Grygorenko OO, Radchenko DS, Dziuba I, Chuprina A, Gubina KE, Moroz YS. Generating Multibillion Chemical Space of Readily Accessible Screening Compounds. iScience. 2020; 23:101681.

9. Gorgulla C, Boeszoermenyi A, Wang ZF, Fischer PD, Coote PW, Padmanabha Das KM, Malets YS, Radchenko DS, Moroz YS, Scott DA, Fackeldey K, Hoffmann M, Iavniuk I, Wagner G, Arthanari H. An open-source drug discovery platform enables ultra-large virtual screens. Nature. 2020; 580:663–8.

10. Acharya A, Agarwal R, Baker MB, Baudry J, Bhowmik D, Boehm S, Byler KG, Chen SY, Coates L, Cooper CJ, Demerdash O, Daidone I, Eblen JD, Ellingson S, Forli S, Glaser J, Gumbart JC, Gunnels J, Hernandez O, Irle S, Kneller DW, Kovalevsky A, Larkin J, Lawrence TJ, LeGrand S, Liu SH, Mitchell JC, Park G, Parks JM, Pavlova A, Petridis L, Poole D, Pouchard L, Ramanathan A, Rogers DM, Santos-Martins D, Scheinberg A, Sedova A, Shen Y, Smith JC, Smith MD, Soto C, Tsaris A, Thavappiragasam M, Tillack AF, Vermaas JV, Vuong VQ, Yin J, Yoo S, Zahran M, Zanetti-Polzi L. Supercomputer-Based Ensemble Docking Drug Discovery Pipeline with Application to Covid-19. J Chem Inf Model. 2020; 60:5832–52.

11. Cherkasov A, Ban F, Li Y, Fallahi M, Hammond GL. Progressive docking: a hybrid QSAR/docking approach for accelerating in silico high throughput screening. J Med Chem. 2006; 49:7466–78.

12. Svensson F, Norinder U, Bender A. Improving Screening Efficiency through Iterative Screening Using Docking and Conformal Prediction. J Chem Inf Model. 2017; 57:439–44.

13. Adeshina YO, Deeds EJ, Karanicolas J. Machine learning classification can reduce false positives in structure-based virtual screening. Proc Natl Acad Sci U S A. 2020; 117:18477–88.

14. Ghose AK, Herbertz T, Pippin DA, Salvino JM, Mallamo JP. Knowledge based prediction of ligand binding modes and rational inhibitor design for kinase drug discovery. J Med Chem. 2008; 51:5149–71.

15. Miduturu CV, Deng X, Kwiatkowski N, Yang W, Brault L, Filippakopoulos P, Chung E, Yang Q, Schwaller J, Knapp S, King RW, Lee JD, Herrgard S, Zarrinkar P, Gray NS. High-throughput kinase profiling: a more efficient approach toward the discovery of new kinase inhibitors. Chem Biol. 2011; 18:868–79.

16. Akritopoulou-Zanze I, Hajduk PJ. Kinase-targeted libraries: the design and synthesis of novel, potent, and selective kinase inhibitors. Drug Discov Today. 2009; 14:291–7.

17. Chen H, Zhou X, Wang A, Zheng Y, Gao Y, Zhou J. Evolutions in fragment-based drug design: the deconstruction-reconstruction approach. Drug Discov Today. 2015; 20:105–13.

18. Pallesen JS, Narayanan D, Tran KT, Solbak SMO, Marseglia G, Sorensen LME, Hoj LJ, Munafo F, Carmona RMC, Garcia AD, Desu HL, Brambilla R, Johansen TN, Popowicz GM, Sattler M, Gajhede M, Bach A. Deconstructing Noncovalent Kelch-like ECH-Associated Protein 1 (Keap1) Inhibitors into Fragments to Reconstruct New Potent Compounds. J Med Chem. 2021.

19. Lamoree B, Hubbard RE. Current perspectives in fragment-based lead discovery (FBLD). Essays Biochem. 2017; 61:453–64.

20. Kirsch P, Hartman AM, Hirsch AKH, Empting M. Concepts and Core Principles of Fragment-Based Drug Design. Molecules. 2019; 24.

21. Bhullar KS, Lagaron NO, McGowan EM, Parmar I, Jha A, Hubbard BP, Rupasinghe HPV. Kinase-targeted cancer therapies: progress, challenges and future directions. Mol Cancer. 2018; 17:48.

22. Noble ME, Endicott JA, Johnson LN. Protein kinase inhibitors: insights into drug design from structure. Science. 2004; 303:1800–5.

23. Taylor SS, Kornev AP. Protein kinases: evolution of dynamic regulatory proteins. Trends Biochem Sci. 2011; 36:65–77.

24. Modi V, Dunbrack RL, Jr. Defining a new nomenclature for the structures of active and inactive kinases. Proc Natl Acad Sci U S A. 2019; 116:6818–27.

25. Zhang H, Pandey S, Travers M, Sun H, Morton G, Madzo J, Chung W, Khowsathit J, Perez-Leal O, Barrero CA, Merali C, Okamoto Y, Sato T, Pan J, Garriga J, Bhanu NV, Simithy J, Patel B, Huang J, Raynal NJ, Garcia BA, Jacobson MA, Kadoch C, Merali S, Zhang Y, Childers W, Abou-Gharbia M, Karanicolas J, Baylin SB, Zahnow CA, Jelinek J, Grana X, Issa JJ. Targeting CDK9 Reactivates Epigenetically Silenced Genes in Cancer. Cell. 2018; 175:1244–58 e26.

26. Kirubakaran P, Morton G, Zhang P, Zhang H, Gordon J, Abou-Gharbia M, Issa J-PJ, Wu J, Childers W, Karanicolas J. Comparative Modeling of CDK9 Inhibitors to Explore Selectivity and Structure-Activity Relationships. bioRxiv. 2020:10.1101/2020.06.08.138602.

27. Gazzard L, Williams K, Chen H, Axford L, Blackwood E, Burton B, Chapman K, Crackett P, Drobnick J, Ellwood C, Epler J, Flagella M, Gancia E, Gill M, Goodacre S, Halladay J, Hewitt J, Hunt H, Kintz S, Lyssikatos J, Macleod C, Major S, Medard G, Narukulla R, Ramiscal J, Schmidt S, Seward E, Wiesmann C, Wu P, Yee S, Yen I, Malek S. Mitigation of Acetylcholine Esterase Activity in the 1,7-Diazacarbazole Series of Inhibitors of Checkpoint Kinase 1. J Med Chem. 2015; 58:5053–74.

28. Bramson HN, Corona J, Davis ST, Dickerson SH, Edelstein M, Frye SV, Gampe RT, Jr., Harris PA, Hassell A, Holmes WD, Hunter RN, Lackey KE, Lovejoy B, Luzzio MJ, Montana V, Rocque WJ, Rusnak D, Shewchuk L, Veal JM, Walker DH, Kuyper LF. Oxindole-based inhibitors of cyclin-dependent kinase 2 (CDK2): design, synthesis, enzymatic activities, and X-ray crystallographic analysis. J Med Chem. 2001; 44:4339–58.

29. Engelman JA, Zejnullahu K, Gale CM, Lifshits E, Gonzales AJ, Shimamura T, Zhao F, Vincent PW, Naumov GN, Bradner JE, Althaus IW, Gandhi L, Shapiro GI, Nelson JM, Heymach JV, Meyerson M, Wong KK, Janne PA. PF00299804, an irreversible pan-ERBB inhibitor, is effective in lung cancer models with EGFR and ERBB2 mutations that are resistant to gefitinib. Cancer Res. 2007; 67:11924–32.

30. Gajiwala KS, Feng J, Ferre R, Ryan K, Brodsky O, Weinrich S, Kath JC, Stewart A. Insights into the aberrant activity of mutant EGFR kinase domain and drug recognition. Structure. 2013; 21:209–19.

31. Ni ZJ, Barsanti P, Brammeier N, Diebes A, Poon DJ, Ng S, Pecchi S, Pfister K, Renhowe PA, Ramurthy S, Wagman AS, Bussiere DE, Le V, Zhou Y, Jansen JM, Ma S, Gesner TG. 4-(Aminoalkylamino)-3-benzimidazole-quinolinones as potent CHK-1 inhibitors. Bioorg Med Chem Lett. 2006; 16:3121–4.

32. Kopecky DJ, Hao X, Chen Y, Fu J, Jiao X, Jaen JC, Cardozo MG, Liu J, Wang Z, Walker NP, Wesche H, Li S, Farrelly E, Xiao SH, Kayser F. Identification and optimization of N3,N6-diaryl-1H-pyrazolo[3,4-d]pyrimidine-3,6-diamines as a novel class of ACK1 inhibitors. Bioorg Med Chem Lett. 2008; 18:6352–6.

33. Coley CW, Rogers L, Green WH, Jensen KF. Computer-Assisted Retrosynthesis Based on Molecular Similarity. ACS Cent Sci. 2017; 3:1237–45.

34. Liu B, Ramsundar B, Kawthekar P, Shi J, Gomes J, Luu Nguyen Q, Ho S, Sloane J, Wender P, Pande V. Retrosynthetic Reaction Prediction Using Neural Sequence-to-Sequence Models. ACS Cent Sci. 2017; 3:1103–13.

35. Coley CW, Green WH, Jensen KF. Machine Learning in Computer-Aided Synthesis Planning. Acc Chem Res. 2018; 51:1281–9.

36. Lee AA, Yang Q, Sresht V, Bolgar P, Hou X, Klug-McLeod JL, Butler CR. Molecular Transformer unifies reaction prediction and retrosynthesis across pharma chemical space. Chem Commun (Camb). 2019; 55:12152–5.

37. Genheden S, Thakkar A, Chadimova V, Reymond JL, Engkvist O, Bjerrum E. AiZynthFinder: a fast, robust and flexible open-source software for retrosynthetic planning. J Cheminform. 2020; 12:70.

38. Shibukawa R, Ishida S, Yoshizoe K, Wasa K, Takasu K, Okuno Y, Terayama K, Tsuda K. CompRet: a comprehensive recommendation framework for chemical synthesis planning with algorithmic enumeration. J Cheminform. 2020; 12:52.

39. Lin K, Xu Y, Pei J, Lai L. Automatic retrosynthetic route planning using template-free models. Chemical Science. 2020; 11:3355–64.

40. Daylight Theory: SMARTS - A Language for Describing Molecular Patterns. 2020 [updated 2020; cited 2020 Feb 27, 2020]; Available from: https://www.daylight.com/dayhtml/doc/theory/theory.smarts.html.

41. RDKit: Open-source cheminformatics (www.rdkit.org).

42. Bollag G, Hirth P, Tsai J, Zhang J, Ibrahim PN, Cho H, Spevak W, Zhang C, Zhang Y, Habets G, Burton EA, Wong B, Tsang G, West BL, Powell B, Shellooe R, Marimuthu A, Nguyen H, Zhang KY, Artis DR, Schlessinger J, Su F, Higgins B, Iyer R, D’Andrea K, Koehler A, Stumm M, Lin PS, Lee RJ, Grippo J, Puzanov I, Kim KB, Ribas A, McArthur GA, Sosman JA, Chapman PB, Flaherty KT, Xu X, Nathanson KL, Nolop K. Clinical efficacy of a RAF inhibitor needs broad target blockade in BRAF-mutant melanoma. Nature. 2010; 467:596–9.

43. Tsai J, Lee JT, Wang W, Zhang J, Cho H, Mamo S, Bremer R, Gillette S, Kong J, Haass NK, Sproesser K, Li L, Smalley KS, Fong D, Zhu YL, Marimuthu A, Nguyen H, Lam B, Liu J, Cheung I, Rice J, Suzuki Y, Luu C, Settachatgul C, Shellooe R, Cantwell J, Kim SH, Schlessinger J, Zhang KY, West BL, Powell B, Habets G, Zhang C, Ibrahim PN, Hirth P, Artis DR, Herlyn M, Bollag G. Discovery of a selective inhibitor of oncogenic B-Raf kinase with potent antimelanoma activity. Proc Natl Acad Sci U S A. 2008; 105:3041–6.

44. Steinbrecher TB, Dahlgren M, Cappel D, Lin T, Wang L, Krilov G, Abel R, Friesner R, Sherman W. Accurate Binding Free Energy Predictions in Fragment Optimization. J Chem Inf Model. 2015; 55:2411–20.

45. Durrant JD, Amaro RE, McCammon JA. AutoGrow: a novel algorithm for protein inhibitor design. Chem Biol Drug Des. 2009; 73:168–78.

46. Hoffer L, Renaud JP, Horvath D. In silico fragment-based drug discovery: setup and validation of a fragment-to-lead computational protocol using S4MPLE. J Chem Inf Model. 2013; 53:836–51.

47. Hawkins PC, Skillman AG, Warren GL, Ellingson BA, Stahl MT. Conformer generation with OMEGA: algorithm and validation using high quality structures from the Protein Databank and Cambridge Structural Database. J Chem Inf Model. 2010; 50:572–84.

48. Park H, Zhou G, Baek M, Baker D, DiMaio F. Force Field Optimization Guided by Small Molecule Crystal Lattice Data Enables Consistent Sub-Angstrom Protein-Ligand Docking. J Chem Theory Comput. 2021; 17:2000–10.

49. Alford RF, Leaver-Fay A, Jeliazkov JR, O’Meara MJ, DiMaio FP, Park H, Shapovalov MV, Renfrew PD, Mulligan VK, Kappel K, Labonte JW, Pacella MS, Bonneau R, Bradley P, Dunbrack RL, Jr., Das R, Baker D, Kuhlman B, Kortemme T, Gray JJ. The Rosetta All-Atom Energy Function for Macromolecular Modeling and Design. J Chem Theory Comput. 2017; 13:3031–48.

50. Jencks WP. On the attribution and additivity of binding energies. Proc Natl Acad Sci U S A. 1981; 78:4046–50.

51. Barelier S, Cummings JA, Rauwerdink AM, Hitchcock DS, Farelli JD, Almo SC, Raushel FM, Allen KN, Shoichet BK. Substrate deconstruction and the nonadditivity of enzyme recognition. J Am Chem Soc. 2014; 136:7374–82.

52. Kramer C, Fuchs JE, Liedl KR. Strong nonadditivity as a key structure-activity relationship feature: distinguishing structural changes from assay artifacts. J Chem Inf Model. 2015; 55:483–94.

53. Nasief NN, Hangauer D. Additivity or cooperativity: which model can predict the influence of simultaneous incorporation of two or more functionalities in a ligand molecule? Eur J Med Chem. 2015; 90:897–915.

54. Cockroft SL, Hunter CA. Chemical double-mutant cycles: dissecting non-covalent interactions. Chem Soc Rev. 2007; 36:172–88.

55. Sydow D, Schmiel P, Mortier J, Volkamer A. KinFragLib: Exploring the Kinase Inhibitor Space Using Subpocket-Focused Fragmentation and Recombination. J Chem Inf Model. 2020.

56. Zhang L, Yang Y, Zhou H, Zheng Q, Li Y, Zheng S, Zhao S, Chen D, Fan C. Structure-activity study of quinazoline derivatives leading to the discovery of potent EGFR-T790M inhibitors. Eur J Med Chem. 2015; 102:445–63.

57. Das D, Hong J. Recent advancements of 4-aminoquinazoline derivatives as kinase inhibitors and their applications in medicinal chemistry. Eur J Med Chem. 2019; 170:55–72.

58. Cavalluzzi MM, Mangiatordi GF, Nicolotti O, Lentini G. Ligand efficiency metrics in drug discovery: the pros and cons from a practical perspective. Expert Opin Drug Discov. 2017; 12:1087–104.

59. Larsson A, Jansson A, Aberg A, Nordlund P. Efficiency of hit generation and structural characterization in fragment-based ligand discovery. Curr Opin Chem Biol. 2011; 15:482–8.

60. Hopkins AL, Keseru GM, Leeson PD, Rees DC, Reynolds CH. The role of ligand efficiency metrics in drug discovery. Nat Rev Drug Discov. 2014; 13:105–21.

61. Ke YY, Coumar MS, Shiao HY, Wang WC, Chen CW, Song JS, Chen CH, Lin WH, Wu SH, Hsu JT, Chang CM, Hsieh HP. Ligand efficiency based approach for efficient virtual screening of compound libraries. Eur J Med Chem. 2014; 83:226–35.

62. Ballester PJ. Selecting machine-learning scoring functions for structure-based virtual screening. Drug Discov Today Technol. 2019; 32–33:81-7.

63. Chaput L, Martinez-Sanz J, Saettel N, Mouawad L. Benchmark of four popular virtual screening programs: construction of the active/decoy dataset remains a major determinant of measured performance. J Cheminform. 2016; 8:56.

64. Lagarde N, Zagury JF, Montes M. Benchmarking Data Sets for the Evaluation of Virtual Ligand Screening Methods: Review and Perspectives. J Chem Inf Model. 2015; 55:1297–307.

65. Reau M, Langenfeld F, Zagury JF, Lagarde N, Montes M. Decoys Selection in Benchmarking Datasets: Overview and Perspectives. Front Pharmacol. 2018; 9:11.

66. Stein RM, Yang Y, Balius TE, O’Meara MJ, Lyu J, Young J, Tang K, Shoichet BK, Irwin JJ. Property-Unmatched Decoys in Docking Benchmarks. J Chem Inf Model. 2021; 61:699–714.

67. Yang Y, Zhang Y, Hua Y, Chen X, Fan Y, Wang Y, Liang L, Deng C, Lu T, Chen Y, Liu H. In Silico Design and Analysis of a Kinase-Focused Combinatorial Library Considering Diversity and Quality. J Chem Inf Model. 2020; 60:92–107.

68. Sydow D, Schmiel P, Mortier J, Volkamer A. KinFragLib: Exploring the Kinase Inhibitor Space Using Subpocket-Focused Fragmentation and Recombination. J Chem Inf Model. 2020; 60:6081–94.

69. Schneider G, Fechner U. Computer-based de novo design of drug-like molecules. Nat Rev Drug Discov. 2005; 4:649–63.

70. Hoffer L, Muller C, Roche P, Morelli X. Chemistry-driven Hit-to-lead Optimization Guided by Structure-based Approaches. Mol Inform. 2018; 37:e1800059.

71. Shan J, Pan X, Wang X, Xiao X, Ji C. FragRep: A Web Server for Structure-Based Drug Design by Fragment Replacement. J Chem Inf Model. 2020; 60:5900–6.

72. Metz A, Wollenhaupt J, Glockner S, Messini N, Huber S, Barthel T, Merabet A, Gerber HD, Heine A, Klebe G, Weiss MS. Frag4Lead: growing crystallographic fragment hits by catalog using fragment-guided template docking. Acta Crystallogr D Struct Biol. 2021; 77:1168–82.

73. Spiegel JO, Durrant JD. AutoGrow4: an open-source genetic algorithm for de novo drug design and lead optimization. J Cheminform. 2020; 12:25.

74. Green H, Koes DR, Durrant JD. DeepFrag: a deep convolutional neural network for fragment-based lead optimization. Chemical Science. 2021.

75. Hoffer L, Voitovich YV, Raux B, Carrasco K, Muller C, Fedorov AY, Derviaux C, Amouric A, Betzi S, Horvath D, Varnek A, Collette Y, Combes S, Roche P, Morelli X. Integrated Strategy for Lead Optimization Based on Fragment Growing: The Diversity-Oriented-Target-Focused-Synthesis Approach. J Med Chem. 2018; 61:5719–32.

76. Konze KD, Bos PH, Dahlgren MK, Leswing K, Tubert-Brohman I, Bortolato A, Robbason B, Abel R, Bhat S. Reaction-Based Enumeration, Active Learning, and Free Energy Calculations To Rapidly Explore Synthetically Tractable Chemical Space and Optimize Potency of Cyclin-Dependent Kinase 2 Inhibitors. J Chem Inf Model. 2019; 59:3782–93.

77. Ghanakota P, Bos PH, Konze KD, Staker J, Marques G, Marshall K, Leswing K, Abel R, Bhat S. Combining Cloud-Based Free-Energy Calculations, Synthetically Aware Enumerations, and Goal-Directed Generative Machine Learning for Rapid Large-Scale Chemical Exploration and Optimization. J Chem Inf Model. 2020; 60:4311–25.

78. Gentile F, Agrawal V, Hsing M, Ton AT, Ban F, Norinder U, Gleave ME, Cherkasov A. Deep Docking: A Deep Learning Platform for Augmentation of Structure Based Drug Discovery. ACS Cent Sci. 2020; 6:939–49.

79. Yang Y, Yao K, Repasky MP, Leswing K, Abel R, Shoichet B, Jerome SV. Efficient Exploration of Chemical Space with Docking and Deep-Learning. chemRxiv. 2021; 10.26434/chemrxiv.14153819.v1.

80. Free SM, Jr., Wilson JW. A Mathematical Contribution to Structure-Activity Studies. J Med Chem. 1964; 7:395–9.

81. Baum B, Muley L, Smolinski M, Heine A, Hangauer D, Klebe G. Non-additivity of functional group contributions in protein-ligand binding: a comprehensive study by crystallography and isothermal titration calorimetry. J Mol Biol. 2010; 397:1042–54.

82. Gu S, Smith MS, Yang Y, Irwin JJ, Shoichet BK. Ligand Strain Energy in Large Library Docking. J Chem Inf Model. 2021.

83. Zhou H, Cao H, Skolnick J. FRAGSITE: A Fragment-Based Approach for Virtual Ligand Screening. J Chem Inf Model. 2021; 61:2074–89.

84. Ustach VD, Lakkaraju SK, Jo S, Yu W, Jiang W, MacKerell AD, Jr. Optimization and Evaluation of Site-Identification by Ligand Competitive Saturation (SILCS) as a Tool for Target-Based Ligand Optimization. J Chem Inf Model. 2019; 59:3018–35.

85. Schuller M, Correy GJ, Gahbauer S, Fearon D, Wu T, Diaz RE, Young ID, Carvalho Martins L, Smith DH, Schulze-Gahmen U, Owens TW, Deshpande I, Merz GE, Thwin AC, Biel JT, Peters JK, Moritz M, Herrera N, Kratochvil HT, Consortium QSB, Aimon A, Bennett JM, Brandao Neto J, Cohen AE, Dias A, Douangamath A, Dunnett L, Fedorov O, Ferla MP, Fuchs MR, Gorrie-Stone TJ, Holton JM, Johnson MG, Krojer T, Meigs G, Powell AJ, Rack JGM, Rangel VL, Russi S, Skyner RE, Smith CA, Soares AS, Wierman JL, Zhu K, O’Brien P, Jura N, Ashworth A, Irwin JJ, Thompson MC, Gestwicki JE, von Delft F, Shoichet BK, Fraser JS, Ahel I. Fragment binding to the Nsp3 macrodomain of SARS-CoV-2 identified through crystallographic screening and computational docking. Sci Adv. 2021; 7.

86. Wang L, Chambers J, Abel R. Protein-Ligand Binding Free Energy Calculations with FEP. Methods Mol Biol. 2019; 2022:201–32.

87. Lombardo F, Desai PV, Arimoto R, Desino KE, Fischer H, Keefer CE, Petersson C, Winiwarter S, Broccatelli F. In Silico Absorption, Distribution, Metabolism, Excretion, and Pharmacokinetics (ADME-PK): Utility and Best Practices. An Industry Perspective from the International Consortium for Innovation through Quality in Pharmaceutical Development. J Med Chem. 2017; 60:9097–113.

88. Fabbro D, Cowan-Jacob SW, Moebitz H. Ten things you should know about protein kinases: IUPHAR Review 14. Br J Pharmacol. 2015; 172:2675–700.

89. Waterhouse A, Bertoni M, Bienert S, Studer G, Tauriello G, Gumienny R, Heer FT, de Beer TAP, Rempfer C, Bordoli L, Lepore R, Schwede T. SWISS-MODEL: homology modelling of protein structures and complexes. Nucleic Acids Res. 2018; 46:W296–W303.

