## Supplemental Figures and Tables for "Efficient Hit-to-Lead Searching of Kinase Inhibitor Chemical Space via Computational Fragment Merging"

### Supporting Figures

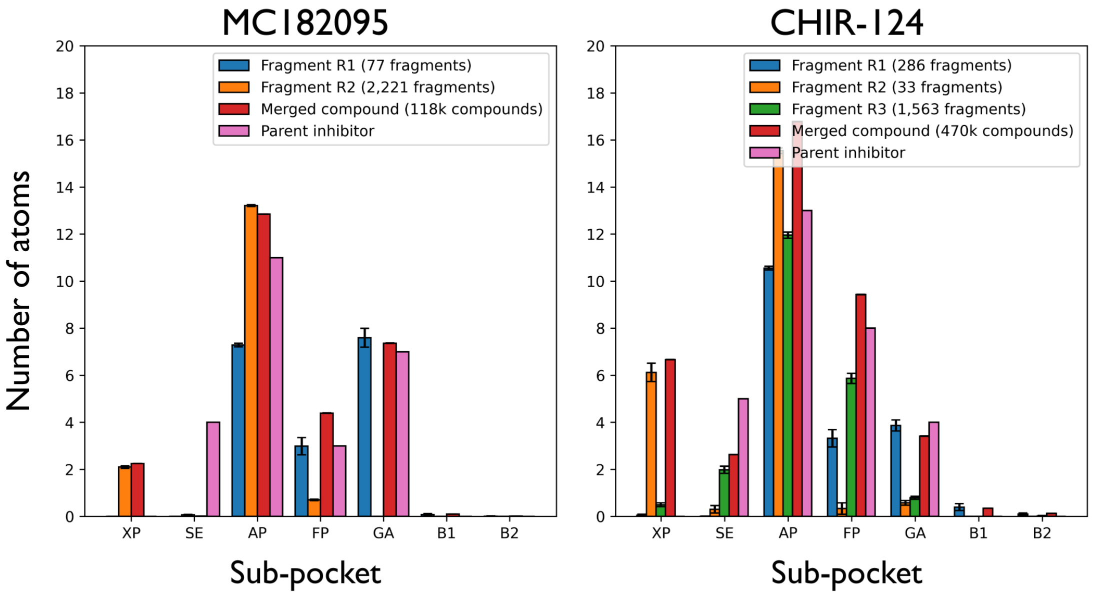


**Figure S1: Merged inhibitors occupy the same sub-pockets of the binding site as their component fragments.** For each ligand, we assigned each atom to the closest KinFragLib-defined sub-pockets within the kinase active site (AP=adenine-binding region, SE=solvent-exposed region, FP=front pocket, GA=gate area, B1=back pocket I, B2=back pocket II, and XP=any remaining ligand atoms). From the crystal structure of MC180295 bound to CDK9 (*pink*), for example, the inhibitor has a total of 25 non-hydrogen atoms. Of these, 11 atoms reside in the AP sub-pocket, 7 in the GA sub-pocket, 4 in the SE sub-pocket, and 3 in the FP sub-pocket. Among the R1 fragments comprising this library (*blue*), they typically fill the GA and FP sub-pockets to a similar extent as MC180295 itself, and partially fill the AP sub-pocket (the latter is occupied by the shared hinge-binding core). The R2 fragments (*orange*) typically fill the remainder of the AP sub-pocket, and occasionally have substituents that extend slightly outside this region (and are thus designated XP). The merged fragments (*red*) exhibit a pattern corresponding to the union of the two fragment libraries: essentially a sum of the non-overlapping regions, but less in the AP sub-pocket because this region harbors the hinge-binding core that is shared between the fragments. Error bars correspond to SEM for each of the fragment libraries / merged inhibitors. No error bars are shown for the parent inhibitor, because these counts come from a single crystal structure in each case.

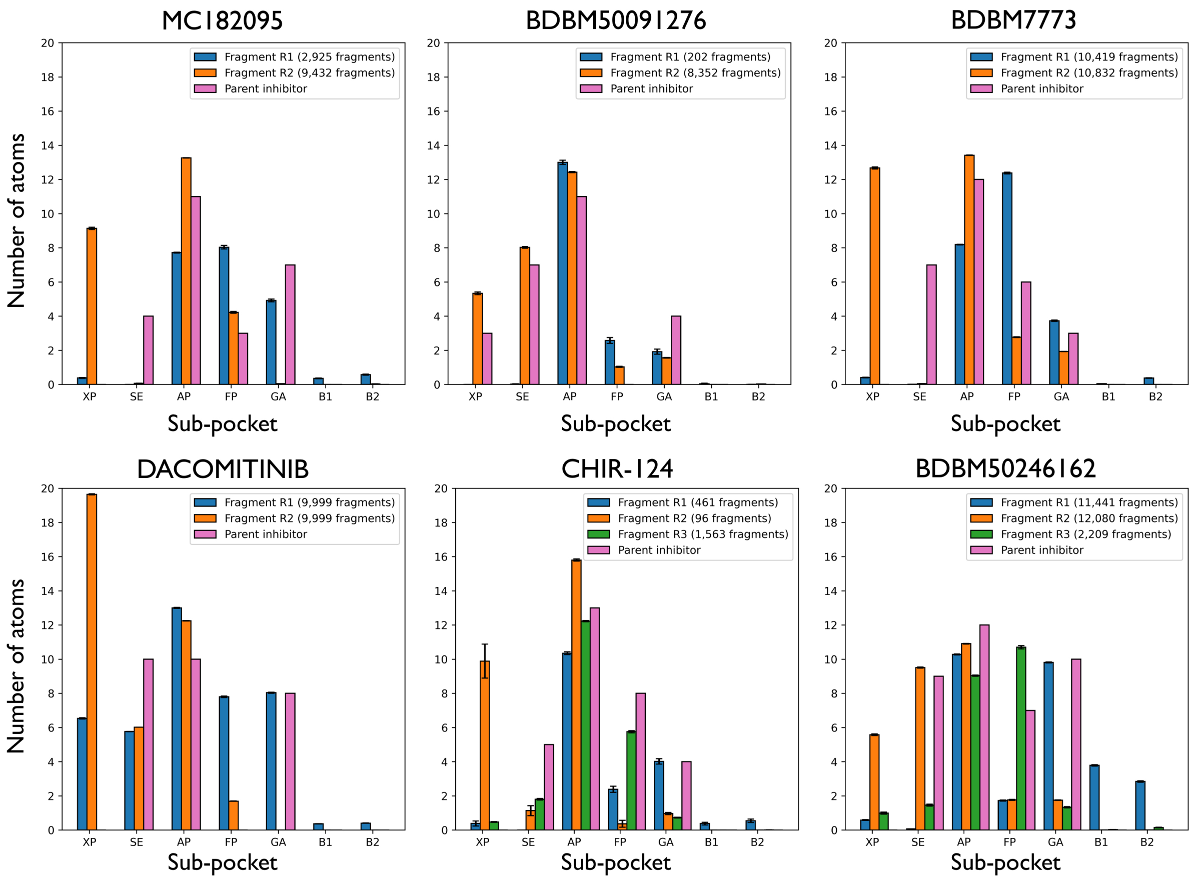


**Figure S2: Fragment libraries occupy the similar sub-pockets as their parent inhibitors.** For each of the building blocks used in assembling the large screening libraries (**Table 1**), we assigned each atom to the closest KinFragLib-defined sub-pockets within the kinase active site. Analysis of the merged inhibitors for this large library as not included (as they were in **Figure S1**), because our method does not necessitate explicit model-building for very large libraries. For each library, at least one of the fragments typically occupies the same pockets as the parent inhibitor (*pink*) with a similar number of atoms. Occupancy of the hinge-binding region (AP), front pocket (FP), and gatekeeper (GA) regions closely matches those of the known inhibitors; this is reassuring, because these buried sub-pockets are the primary contributors to binding. The main difference these distributions comes through slightly less occupancy at the SE (solvent exposed) sub-pocket relative to the parent inhibitor, which the fragment libraries instead assign to the XP sub-pocket ("other regions"). Inspection of the modeled fragments shows that this difference arises in part from bulky substituents which do not fit in the canonical SE region. Error bars correspond to SEM for each of the fragment libraries / merged inhibitors. No error bars are shown for the parent inhibitor, because these counts come from a single crystal structure in each case.

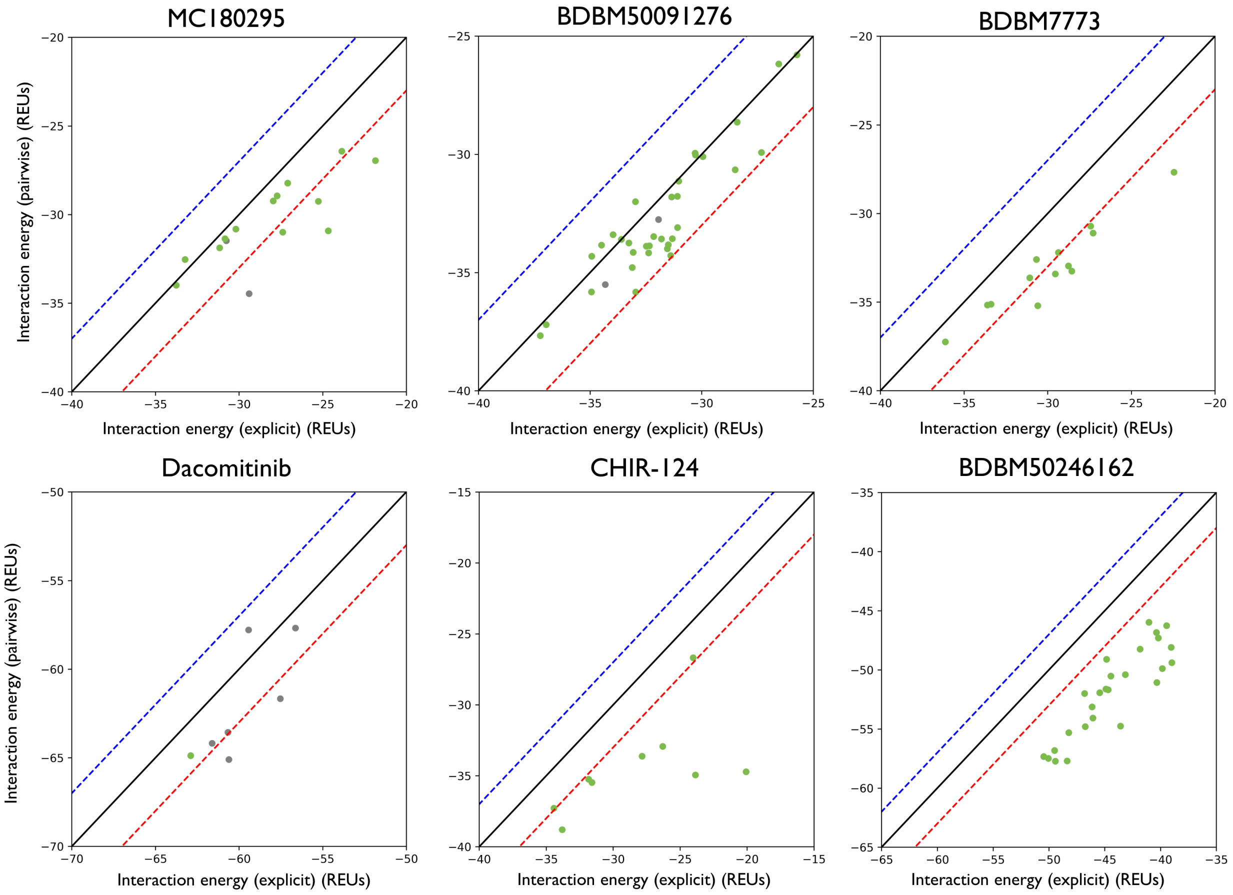


**Figure S3: Pairwise additivity of analogs known to be active.** Interaction energies are reported for each of the analogs with known activity, either estimated from the component fragments using the pairwise approximation (y-axis), or as explicitly calculated from the re-refined model of the complete inhibitor (x-axis). The boundaries for which the additivity approximation holds closely (3 Rosetta energy units in either direction, as used in **Figure 5**) are marked in dashed lines.

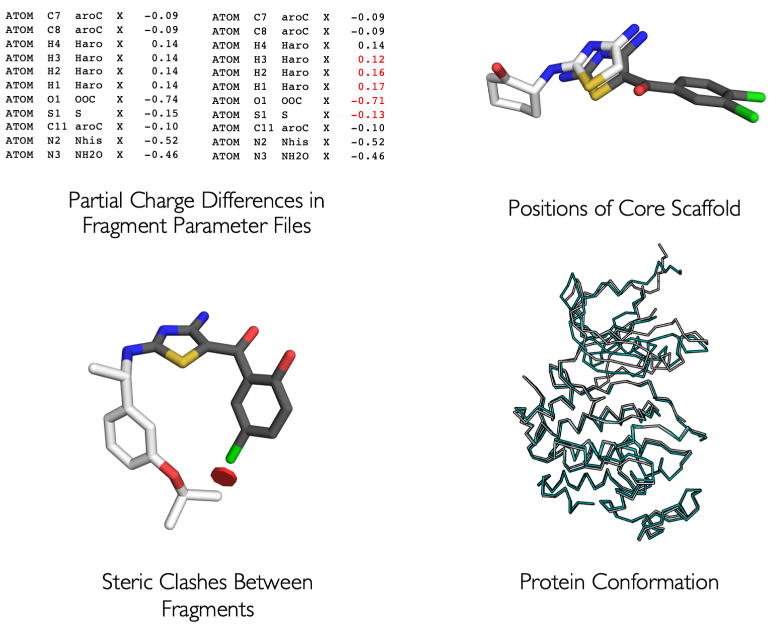


**Figure S4: Basis for non-additivity in modeling kinase inhibitors from fragments.** Early calculations yield results that were not strongly pairwise additive, making these unsuitable for estimating interaction energies of the merged compounds. We found that this non-additivity can arise from 1) inconsistent assignment of partial charges, 2) steric clashes between substituents, 3) incompatible positioning of the core hinge-binding motif, or 4) differences in the modeled conformation of the protein in response to different fragments.

### Supporting Tables

| Library | Internal library name | SMARTS string to search PubChem for suitable building blocks | SMARTS string to generate fragments from the resulting building blocks (i.e., by linking onto the hinge-binding core) |
|---|---|---|---|
| MC180295 | R1 | [H]C([H])(Br)[#6](=O)-[#6]-1=[#6]-[#6]=[#6]-[#6]=[#6]-1 | [#6:4]-[#6:2](=[O:1])-[#6:3]Br>>[#6:4]-[#6:2](=[O:1])-[#6:3]-1=[#6](-[#7])-[#7]=[#6](-[#7])-[#16]-1 |
| | R2 | [H][#7]([H])-* | [#6:1]-[#7H2:2]>>[#6:1]-[#7:2]-[#6]-1=[#7]-[#6](-[#7])=[#6]-[#16]-1 |
| | R1_R2 | | [#6:6]-[#6:2](=[O:1])-[#6:3]Br.[#6:5]-[#7H2:4]>>[#6:5]-[#7:4]-[#6]-1=[#7]-[#6](-[#7])=[#6:3](-[#16]-1)-[#6:2](-[#6:6])=[O:1] |
| BDBM50091276 | R1 | [H][#7]([H])-[#6]-1=[#6](I)-[#6]=[#6]-[#7]=[#6]-1 | [#7:6]-[#6:5]-1=[#6:7]-[#7:1]=[#6:2]-[#6:3]=[#6:4]-1I>>[#7:6]-1-[#6:5]-2=[#6:4](-[#6:3]=[#6:2]-[#7:1]=[#6:7]-2)-[#6]-2=[#6]-1-[#7]=[#6]-[#6]=[#6]-2 |
| | R2 | [H][#8]-[#5](-[*,#1])-[#8][H] | [#8]-[#5](-[#8])-[*:1]>>[*:1]-[#6]-1=[#6]-[#6]-2=[#6](-[#7]-[#6]-3=[#6]-[#7]=[#6]-[#6]=[#6]-2-3)-[#7]=[#6]-1 |
| | R1_R2 | | [#7:2]-[#6:3]-1=[#6:4]-[#7:5]=[#6:6]-[#6:7]=[#6:8]-1I.[#8]-[#5](-[#8])-[*:1]>>[*:1]-[#6]-1=[#6]-[#6]-2=[#6](-[#7:2]-[#6:3]-3=[#6:8]-2-[#6:7]=[#6:6]-[#7:5]=[#6:4]-3)-[#7]=[#6]-1 |
| CHIR-124 | R1 | [H][#7]-1-[#6](=O)-[#8]-[#6](=O)-[#6]-2=[#6]-1-[#6]=[#6]-[#6]=[#6]-2 | [O:6]=[#6:5]-1-[#6:7]~[#6:8]-[#7:1]-[#6:2](=[O:3])-[#8:4]-1>>[O:3]=[#6:2]-1-[#7:1]-[#6:8]~[#6:7]-[#6:5]=[#6:4]-1 |
| | R2 | [H][#7]-1-[#6]-2=[#6](-[#6]=[#6]-[#6]=[#6]-2)-[#7]=[#6]-1C([H])([H])[#6](=O)-[#8]C([H])([H])C([H])([H])[H] | [#6]-[#6]-[#6]-[#8]-[#6:3](=[O:4])-[#6:2]-[*:1]>>[*:1]-[#6:2]-1=[#6]-[#6]-2=[#6]-[#6]=[#6]-[#6]=[#6]-2-[#7]-[#6:3]-1=[O:4] |
| | R1_R2 | | [O:6]=[#6:5]-1-[#6:7]~[#6:8]-[#7:1]-[#6:2](=[O:3])-[#8:4]-1.[#6]-[#6]-[#6]-[#8]-[#6](=O)-[#6]-[*:9]>>[*:9]-[#6:4]-1=[#6:5]-[#6:7]~[#6:8]-[#7:1]-[#6:2]-1=[O:3] |
| BDBM7773 | R1 | [H][#7]([H])-[#6]-1=[#6]-[#6]=[#6]-[#6]=[#6]-1 | [#7:1]-[#6:2]-1=[#6:3]-[#6:4]=[#6:5]-[#6:6]=[#6:7]-1>>[#7]\[#6]=[#6]-1\[#6](=O)-[#7:1]-[#6:2]-2=[#6:3]-[#6:4]=[#6:5]-[#6:6]=[#6:7]-1-2 |
| | R2 | [H][#7]([H])-* | [#7:1]-[*:2]>>[*:2]-[#7:1]\[#6]=[#6]-1/[#6](=O)-[#7]-[#6]-2=[#6]-[#6]=[#6]-[#6]=[#6]-1-2 |
| | R1_R2 | | [#7:1]-[#6:2]-1=[#6:3]-[#6:4]=[#6:5]-[#6:6]=[#6:7]-1.[#7:8]-[*:9]>>[*:9]-[#7:8]\[#6]=[#6]-1/[#6](=O)-[#7:1]- |

| | | | |
|---|---|---|---|
| | | | [#6:2]-2=[#6:3]-[#6:4]=[#6:5]-[#6:6]=[#6:7]-1-2 |
| BDBM50246162 | R1 | [H][#7]([H])-* | [#7:2]-[*:1]>>[#7]-[#6]-1=[#7]-[#6]=[#6]-2-[#6](-[#7:2]-[*:1])=[#7]-[#7]-[#6]-2=[#7]-1 |
| | R2 | [H][#7]([H])-[#7]([H])-* | [#7:1]-[#7:2]-[*:3]>>[#7]-[#6]-1=[#7:1]-[#7:2](-[*:3])-[#6]-2=[#7]-[#6](-[#7])=[#7]-[#6]=[#6]-1-2 |
| | R3 | [H][#7]([H])-* | [#7:2]-[*:1]>>[#7]-[#6]-1=[#7]-[#7]-[#6]-2=[#7]-[#6](-[#7:2]-[*:1])=[#7]-[#6]=[#6]-1-2 |
| | R1_R2_R3 | | [#7:2]-[*:1].[#7:3]-[#7:4]-[*:5].[#7:7]-[*:6]>>[*:1]-[#7:2]-[#6]-1=[#7:3]-[#7:4](-[*:5])-[#6]-2=[#7]-[#6](-[#7:7]-[*:6])=[#7]-[#6]=[#6]-1-2 |

**Table S1:** SMARTS templates for transforming building blocks into fragment libraries. The R1 and R2 SMARTS strings encode the reactions needed to transform building blocks into the corresponding fragments (i.e., by adding the hinge-binding scaffold). The R1_R2 SMARTS strings encode all at once the complete set of reactions needed to transform multiple building blocks into the final inhibitor.

| Option | Value |
|---|---|
| Single or double bonds match aromatic bonds | True |
| Chain bonds in the query may match rings in hits | True |
| Atoms must be of the specified isotope | True |
| Allow match to tautomers of the given structure | True |
| Atoms must match the specified charge | False |
| Rings may not be embedded in a larger system | False |
| Remove any explicit hydrogens before searching | False |

**Table S2:** Search parameters for identifying building blocks in PubChem, using SMARTS matching.

| | | | |
|---|---|---|---|
| AA BLOCKS | Ambinter | Chemhere | Sigma-Aldrich |
| ABI Chem | Aurora Fine Chemicals LLC | ChemTik | SynHet - Synthetic Heterocycles |
| AKos Consulting & Solutions | BLD Pharm | Enamine | TimTec |
| Alichem | ChemBridge | Life Chemicals | Vitas-M Laboratory |
| Ambeed | ChemDiv | MuseChem | |

**Table S3:** Vendors included in our search for available building blocks.

| Library | Number of R1 fragments | Number of R2 fragments | Number of R3 fragments | Enumerated library size |
|---|---|---|---|---|
| MC180295 | 77 | 1,704 | N/A | 131,208 |
| BDBM50091276 | 9 | 2,401 | N/A | 21,609 |
| BDBM7773 | 635 | 647 | N/A | 410,845 |
| Dacomitinib | 470 | 482 | N/A | 226,540 |
| CHIR-124 | 152 | 33 | 151 | 757,416 |
| BDBM50246162 | 145 | 78 | 31 | 350,610 |

**Table S4: Smaller libraries for evaluating pairwise additivity approximation.** Determining the appropriateness of the pairwise additivity approximation requires explicitly evaluating compounds' interaction energies in a non-pairwise way; this precludes the use of huge chemical libraries. For this experiment we therefore selected randomly a subset of the building blocks for each reaction. Enumerating these reactions with fewer building blocks provides much smaller chemical libraries that can be explicitly screened for this experiment.

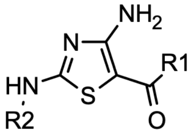


MC180295 core

| Compound name | R1 | R2 | Rank, % |
|---|---|---|---|
| MC180301 | 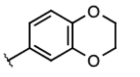 | 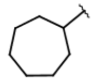 | 0.1 |
| MC180296 | 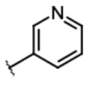 | 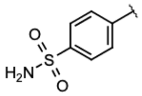 | 1.0 |
| MC180299 | 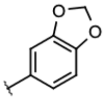 | 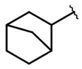 | 1.6 |
| MC180303 | 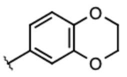 | 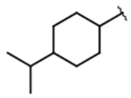 | 2.0 |
| MC180340 | 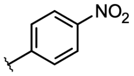 | 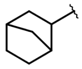 | 3.8 |
| MC180298 | 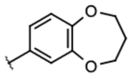 | 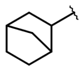 | 4.0 |
| MC180336 | 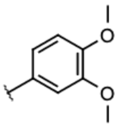 | 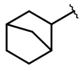 | 4.3 |
| MC180338 | 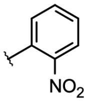 | 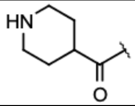 | 5.0 |
| MC180343 | 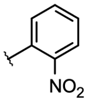 | 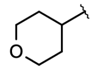 | 5.4 |
| MC180295 | 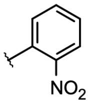 | 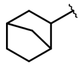 | 12.6 |
| MC180339 | 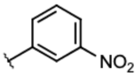 | 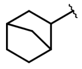 | 15.3 |
| MC180300 | 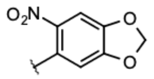 | 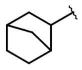 | 16.4 |
| MC180297 | 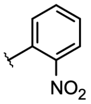 | 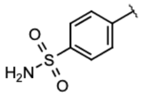 | 17.4 |

| | | | |
|---|---|---|---|
| MC180349 | 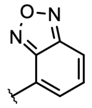 | 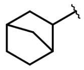 | 23.9 |
| MC180342 | 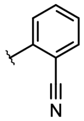 | 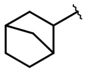 | 44.7 |
| MC180345 | 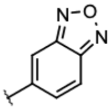 | 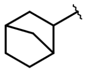 | 45.1 |

**Table S5:** Complete set of analogs of MC180295 previously reported to have activity against CDK9, as used in our study. The rank of each analog in our screen (**Figure 7**) is provided relative to the complete enumerated library used in this experiment (**Table 1**).

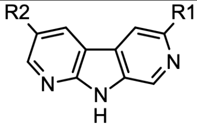


BDBM50091276 core

| Compound name | R1 | R2 | Rank, % |
|---|---|---|---|
| C38 | 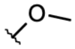 | 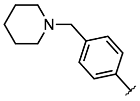 | 0.2 |
| C45 | 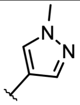 | 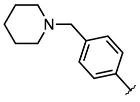 | 0.3 |
| C41 | 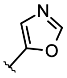 | 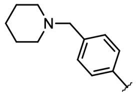 | 0.7 |
| C46 | 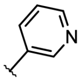 | 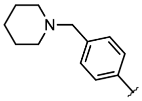 | 0.8 |
| C42 | 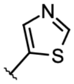 | 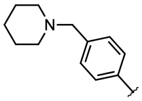 | 0.8 |
| C37 | 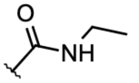 | 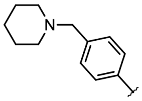 | 0.9 |
| C48 | 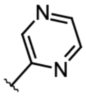 | 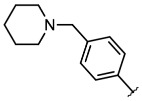 | 1.2 |

| | | | |
|---|---|---|---|
| C43 | 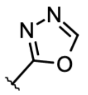 | 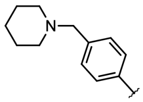 | 1.4 |
| C49 | 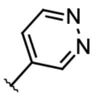 | 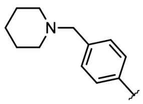 | 2.8 |
| C5 | 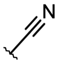 | 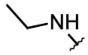 | 3.5 |
| C7 | 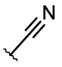 | 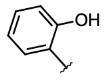 | 4.0 |
| C12 | 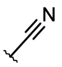 | 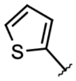 | 4.4 |
| C3 | 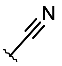 | 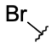 | 5.2 |
| C39 | 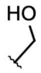 | 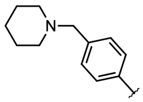 | 6.3 |
| C13 | 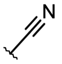 | 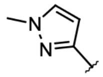 | 6.5 |
| C11 | 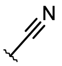 | 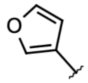 | 6.7 |
| C14 | 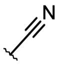 | 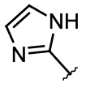 | 7.8 |
| C44 | 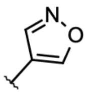 | 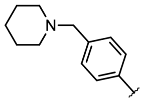 | 9.9 |
| C33 | 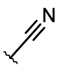 | 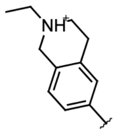 | 10.1 |
| C9 | 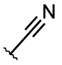 | 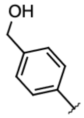 | 12.4 |
| C34 | 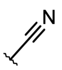 | 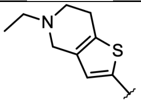 | 12.8 |
| C16 | 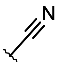 | 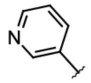 | 13.0 |

| | | | |
|---|---|---|---|
| C15 | 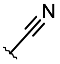 | 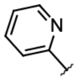 | 13.6 |
| C32 | 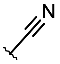 | 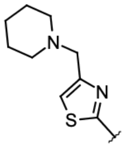 | 16.2 |
| C24 | 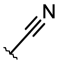 | 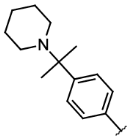 | 20.0 |
| C8 | 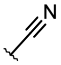 | 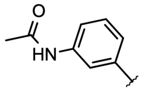 | 20.3 |
| C29 | 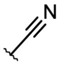 | 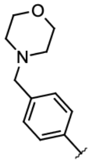 | 20.7 |
| C40 | 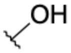 | 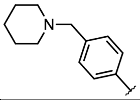 | 20.8 |
| C4 | 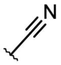 | 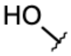 | 21.2 |
| C31 | 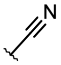 | 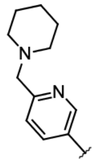 | 21.7 |
| C26 | 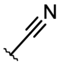 | 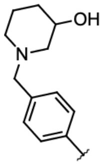 | 22.5 |
| C19 | 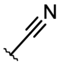 | 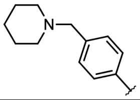 | 23.0 |
| C21 | 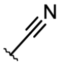 | 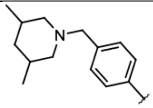 | 23.6 |
| C35 | 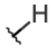 | 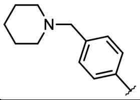 | 24.9 |
| C23 | 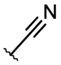 | 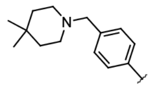 | 26.7 |
| C27 | 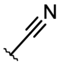 | 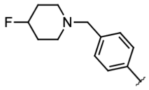 | 27.2 |

| | | | |
|---|---|---|---|
| C25 | 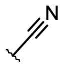 | 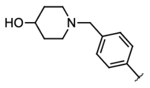 | 28.0 |
| C1 | 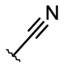 | 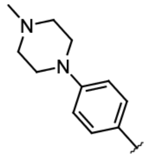 | 29.0 |
| C20 | 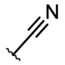 | 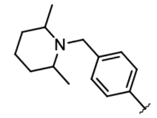 | 29.2 |
| C28 | 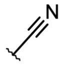 | 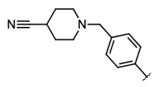 | 29.8 |
| C22 | 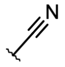 | 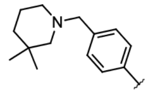 | 30.8 |
| C6 | 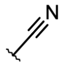 | 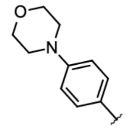 | 31.5 |
| C30 | 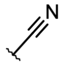 | 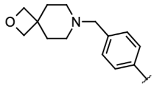 | 32.4 |
| C2 | 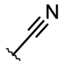 | 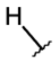 | 81.0 |

**Table S6:** Complete set of analogs of BDBM50091276 previously reported to have activity against CHK1, as used in our study. The rank of each analog in our screen (**Figure 7**) is provided relative to the complete enumerated library used in this experiment (**Table 1**).

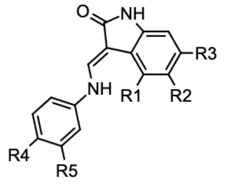


BDBM7773 core

| Compound name | R1 | R2 | R3 | R4 | R5 | Rank, % |
|---|---|---|---|---|---|---|
| C68 | H | H | -CH2-OH | 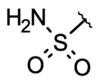 | H | 0.0001 |
| C38 | -OH | H | H | 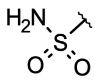 | H | 0.0002 |

| | | | | | | |
|---|---|---|---|---|---|---|
| C18 | H | H | H | 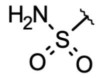 | H | 0.0002 |
| C99 | H | H | H | 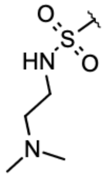 | H | 0.0005 |
| C48 | H | 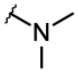 | H | 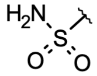 | H | 0.0034 |
| C55 | H | 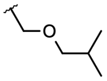 | H | 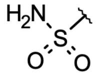 | H | 0.0041 |
| C100 | H | H | H | 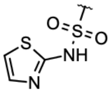 | H | 0.0048 |
| C37 | 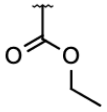 | H | H | 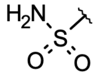 | H | 0.0168 |
| C54 | H | 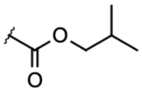 | H | 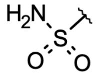 | H | 0.3284 |
| C109 | H | 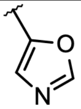 | H | -CH2-S(=O)(=O)-CH2- | | 1.5 |
| C23 | H | 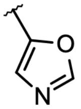 | H | 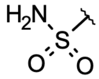 | H | 1.9 |
| C53 | H | 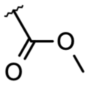 | H | 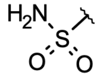 | H | 3.6 |
| C52 | H | 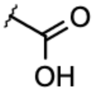 | H | 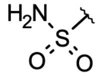 | H | 52.0 |
| C85 | -S-CH=N- | | H | 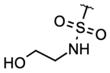 | H | 100.0 |
| C102 | -S-CH=N- | | H | 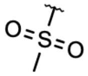 | H | 100.0 |
| C108 | -S-CH=N- | | H | -CH2-S(=O)(=O)-CH2- | | 100.0 |

| | | | | | |
|---|---|---|---|---|---|
| C84 | -S-CH=N- | H | 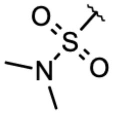 | H | 100.0 |
| C83 | -S-CH=N- | H | 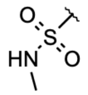 | H | 100.0 |
| C80 | -S-CH=N- | H | 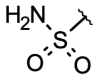 | H | 100.0 |
| C107 | -S-CH=N- | H | 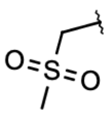 | H | 100.0 |
| C82 | =CH-CH=CH-N= | H | 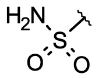 | H | 77.4 |
| C103 | -S-CH=N- | H | 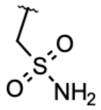 | H | 100.0 |
| C104 | -S-CH=N- | H | 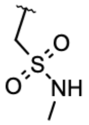 | H | 100.0 |
| C97 | -S-CH=N- | H | 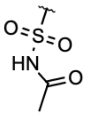 | H | 100.0 |
| C86 | -S-CH=N- | H | 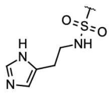 | H | 100.0 |
| C96 | -S-CH=N- | H | 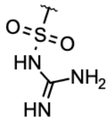 | H | 100.0 |
| C87 | -S-CH=N- | H | 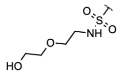 | H | 100.0 |
| C91 | -S-CH=N- | H | 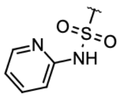 | H | 100.0 |
| C106 | -S-CH=N- | H | 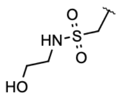 | H | 100.0 |

| | | | | | |
|---|---|---|---|---|---|
| C94 | -S-CH=N- | H | 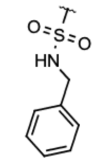 | H | 100.0 |
| C92 | -S-CH=N- | H | 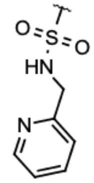 | H | 100.0 |
| C90 | -S-CH=N- | H | 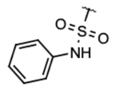 | H | 100.0 |
| C89 | -S-CH=N- | H | 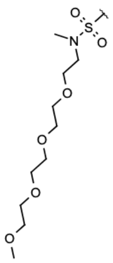 | H | 100.0 |
| C93 | -S-CH=N- | H | 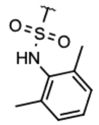 | H | 100.0 |
| C88 | -S-CH=N- | H | 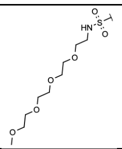 | H | 100.0 |
| C95 | -S-CH=N- | H | 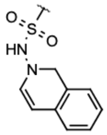 | H | 100.0 |

**Table S7:** Complete set of analogs of BDBM7773 previously reported to have activity against CDK2, as used in our study. The rank of each analog in our screen (**Figure 7**) is provided relative to the complete enumerated library used in this experiment (**Table 1**).

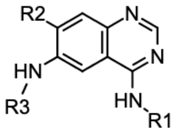


Dacomitinib core

| Compound name | R1 | R2 | R3 | Rank, % |
|---|---|---|---|---|
| C44 | 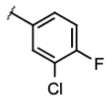 | 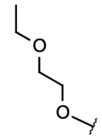 | 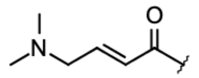 | 7.49E-11 |
| C8H | 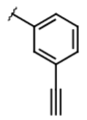 | 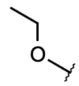 | 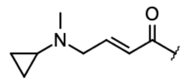 | 6.24E-10 |
| C8E | 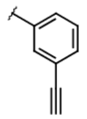 | 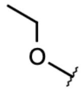 | 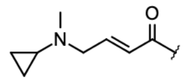 | 4.37E-09 |
| C45 | 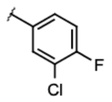 | 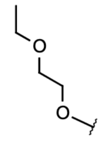 | 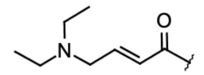 | 1.40E-08 |
| C8G | 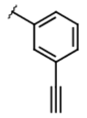 | 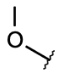 | 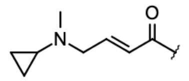 | 5.18E-07 |
| C8I | 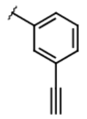 | 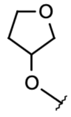 | 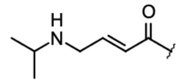 | 7.48E-07 |
| C8D | 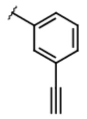 | 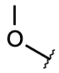 | 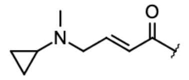 | 3.46E-06 |
| Afatinib | 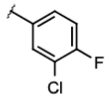 | 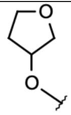 | 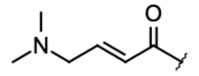 | 4.04E-05 |
| C43 | 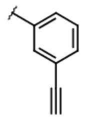 | 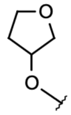 | 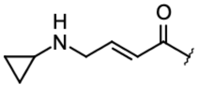 | 5.21E-05 |
| C42 | 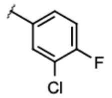 | 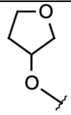 | 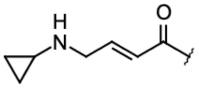 | 1.06E-04 |
| C8F | 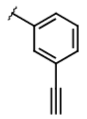 | 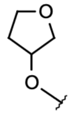 | 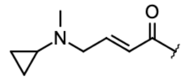 | 3.19E-04 |
| C7B | 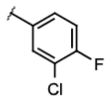 | 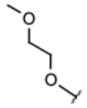 | 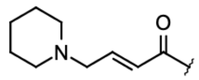 | 3.61E-04 |

| | | | | |
|---|---|---|---|---|
| C8C | 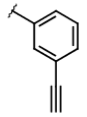 | 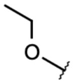 | 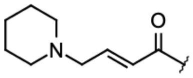 | 2.28E-02 |
| C8B | 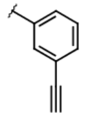 | 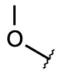 | 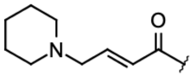 | 99.5 |
| Dacomitinib | 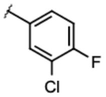 | 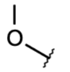 | 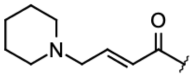 | 99.6 |

**Table S8:** Complete set of analogs of Dacomitinib previously reported to have activity against $EGFR^{T790M}$, as used in our study. The rank of each analog in our screen (**Figure 7**) is provided relative to the complete enumerated library used in this experiment (**Table 1**).

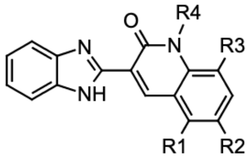


CHIR-124 core

| Compound name | R1 | R2 | R3 | R4 | Rank, % |
|---|---|---|---|---|---|
| C9 | H | Cl | H | 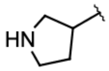 | 4.73E-10 |
| C8 | H | CH3 | H | 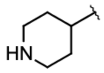 | 1.02E-09 |
| C13 | H | Cl | H | 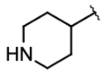 | 2.09E-09 |
| C7 | H | CH3 | H | 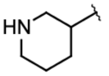 | 5.05E-07 |
| C12 | H | Cl | H | 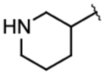 | 9.35E-07 |
| C6 | H | CH3 | H | 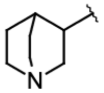 | 2.00E-04 |
| C11 | H | Cl | H | 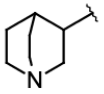 | 3.30E-04 |
| C1 | H | H | H | H | 4.49E-04 |

| | | | | | |
|---|---|---|---|---|---|
| C14 | Cl | Cl | H | 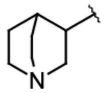  | 22.0 |
| C4 | H | H | H | 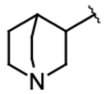  | 26.9 |
| C15 | H | H | CH3 | 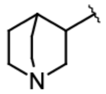  | 100.0 |

**Table S9:** Complete set of analogs of CHIR-124 previously reported to have activity against CHK1, as used in our study. The rank of each analog in our screen (**Figure 7**) is provided relative to the complete enumerated library used in this experiment (**Table 1**).

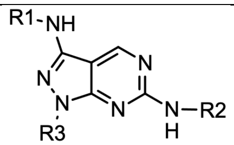


BDBM50246162 core

| **Compound name** | **R1** | **R2** | **R3** | **Rank, %** |
|---|---|---|---|---|
| C33 | 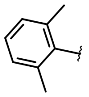 | 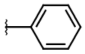 | 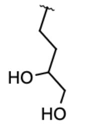  | 7.9E-17 |
| C34 | 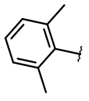 | 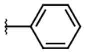 | 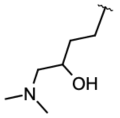  | 1.4E-15 |
| C30 | 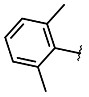 | 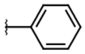 | 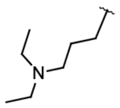  | 3.7E-15 |
| C32 | 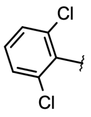  | 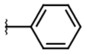 | 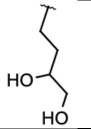  | 1.2E-12 |
| C29 | 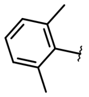 | 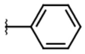 | 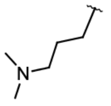  | 2.0E-12 |
| C7 | 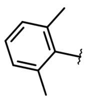 | 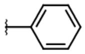 | 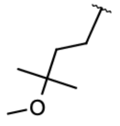  | 2.5E-12 |

| | | | | |
|---|---|---|---|---|
| C31 | 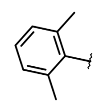 | 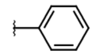 | 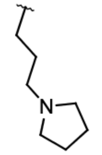 | 2.5E-11 |
| C22 | 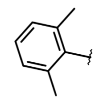 | 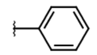 | 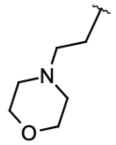 | 4.4E-11 |
| C27 | 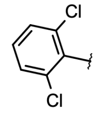 | 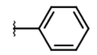 | 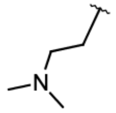 | 7.0E-11 |
| C28 | 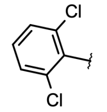 | 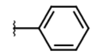 | 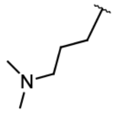 | 7.3E-09 |
| C8 | 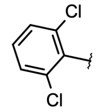 | 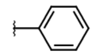 | 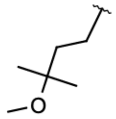 | 8.9E-09 |
| C25 | 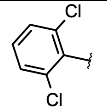 | 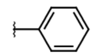 | 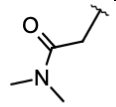 | 2.1E-08 |
| C21 | 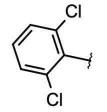 | 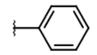 | 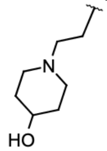 | 2.4E-08 |
| C2 | 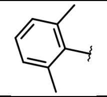 | 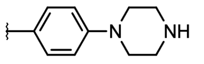 | 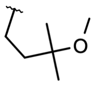 | 5.0E-8 |
| C26 | 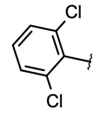 | 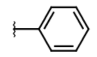 | 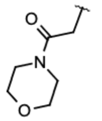 | 9.2E-08 |
| C23 | 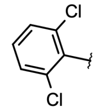 | 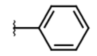 | 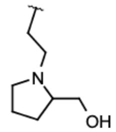 | 1.2E-07 |
| C24 | 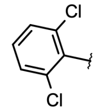 | 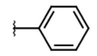 | 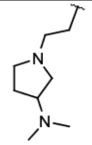 | 2.1E-07 |
| C13 | 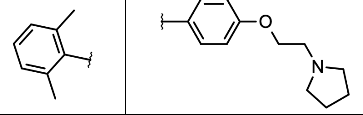 | | 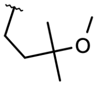 | 2.8E-07 |
| C12 | 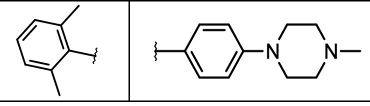 | | 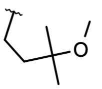 | 1.9E-06 |

| | | | | |
|---|---|---|---|---|
| C20 | 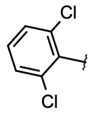 | 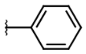 | 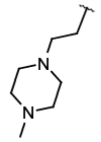 | 6.6E-06 |
| C16 | 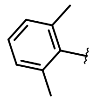 | 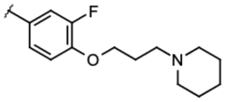 | 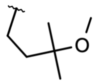 | 7.0E-06 |
| C14 | 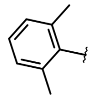 | 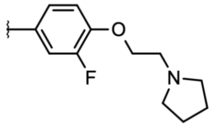 | 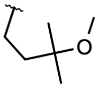 | 9.4E-06 |
| C11 | 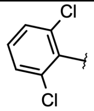 | 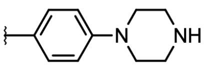 | 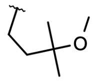 | 3.6E-05 |
| C18 | 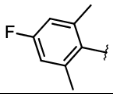 | 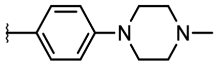 | 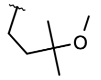 | 1.2E-04 |
| C17 | 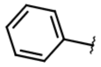 | 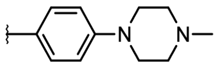 | 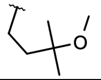 | 2.2E-04 |
| C15 | 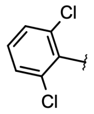 | 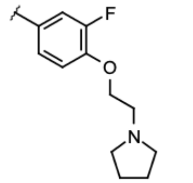 | 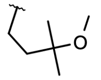 | 2.5E-03 |

**Table S10:** Complete set of analogs of BDBM50246162 previously reported to have activity against ACK1, as used in our study. The rank of each analog in our screen (**Figure 7**) is provided relative to the complete enumerated library used in this experiment (**Table 1**).
